# Retapamulin-assisted ribosome profiling reveals the alternative bacterial proteome

**DOI:** 10.1101/520783

**Authors:** Sezen Meydan, James Marks, Dorota Klepacki, Virag Sharma, Pavel V. Baranov, Andrew E. Firth, Tōnu Margus, Amira Kefi, Nora Vázquez-Laslop, Alexander S. Mankin

**Affiliations:** Center for Biomolecular Sciences - m/c 870, University of Illinois at Chicago, 900 S. Ashland Ave., Chicago, IL 60607, USA; CRTD-DFG Center for Regenerative Therapies Dresden Carl Gustav Carus Faculty of Medicine, Technische Universität Dresden, Dresden; Paul Langerhans Institute Dresden of the Helmholtz Center Munich at University Hospital, Carl Gustav Carus Faculty of Medicine, Technische Universität Dresden, Dresden; German Center for Diabetes Research, Munich, Neuherberg, Germany; School of Biochemistry and Cell Biology, University College Cork, Cork T12 YN60, Ireland; Division of Virology, Department of Pathology, University of Cambridge, Cambridge CB2 1QP, UK

## Abstract

The use of alternative translation initiation sites enables production of more than one protein from a single gene, thereby expanding cellular proteome. Although several such examples have been serendipitously found in bacteria, genome-wide mapping of alternative translation start sites has been unattainable. We found that the antibiotic retapamulin specifically arrests initiating ribosomes at start codons of the genes. Retapamulin-enhanced Ribo-seq analysis (Ribo-RET) not only allowed mapping of conventional initiation sites at the beginning of the genes but, strikingly, it also revealed putative internal start sites in a number of *Escherichia coli* genes. Experiments demonstrated that the internal start codons can be recognized by the ribosomes and direct translation initiation in vitro and in vivo. Proteins, whose synthesis is initiated at an internal in-frame and out-of-frame start sites, can be functionally important and contribute to the ‘alternative’ bacterial proteome. The internal start sites my also play regulatory roles in gene expression.

## INTRODUCTION

A broader diversity of proteins with specialized functions can augment cell reproduction capacity, optimize its metabolism and facilitate survival in the ever-changing environment. However, the fitness gain from acquiring a capacity of making a new protein is counterbalanced with the cost of expanding the size of the genome. This conundrum is particularly onerous in bacteria whose genomes are highly streamlined. Furthermore, expanding the protein repertoire by gene duplication and subsequent specialization could be precarious because similar nucleotide sequences are prone to homologous recombination leading to genome instability (Birch et al., 1991; Koressaar and Remm, 2012).

In order to overcome the genome expansion constraints, bacteria have evolved several strategies for diversifying their proteome without expanding the size of the genome. Some of these strategies, collectively known as recoding, commonly require deviations from the standard rules of translation (Atkins et al., 2016; Baranov et al., 2015). In the recoding scenarios, translation of the nucleotide sequence is initiated at a unique start codon of an open reading frame (ORF), but due to programmed ribosomal frameshifting or stop codon readthrough, a fraction of ribosomes produces a polypeptide whose sequence deviates from that encoded in the main ORF, leading to generation of more than one protein from a single gene.

Another possible way for producing diverse polypeptides from a single ORF and, in this case, without deviating from the canonical rules of decoding, is the utilization of alternative internally located start codons.

While locating the start codon of a protein coding sequence is one of the most critical functions of the ribosome, the rules for translation initiation site (TIS) recognition are fairly flexible. Even though AUG is the most commonly used start codon, other triplets such as GUG, UUG, CUG, AUU and AUC, that differ from AUG by a single nucleotide, can be also decoded by the initiator fMet-tRNA and can direct with varying efficiencies initiation of translation (Villegas and Kropinski, 2008). In bacterial genes, the start codon is often preceded by a Shine-Dalgarno (SD) sequence, which is complementary to the nucleotide sequence at the 3’ end of the 16S rRNA (Shine and Dalgarno, 1975). The presence of a SD sequence, however, is neither sufficient, nor always required for efficient translation initiation (Gualerzi and Pon, 2015; Li et al., 2014; Moll et al., 2002; Woolstenhulme et al., 2015). The mRNA spatial structure (Gualerzi and Pon, 2015; Kozak, 2005) also modulates the efficiency of the start codon recognition, but the extent of its contribution, as well as the role of other yet unknown mRNA features, remain poorly understood (Espah Borujeni et al., 2014; Li et al., 2014; Osterman et al., 2013; Vimberg et al., 2007). The flexible rules of start codon selection should make it difficult for the cell to completely avoid occasional initiation of translation at some internal codons. If this is really the case, certain internal start sites and the resulting protein products could be subject of the evolutionary selection and, thus, could be retained and optimized to eventually benefit the host cell.

Although translation of the majority of the bacterial genes is initiated at a unique TIS, designated herein as primary (pTIS), several examples of genes with an additional internal TIS (iTIS) have been serendipitously discovered, usually during the purification and functional characterization of the primary protein (reviewed in (Meydan et al., 2018)). In these genes, translation initiated at the pTIS results in production of the full-length (primary) protein, while ribosomes which initiate translation at the in-frame iTIS synthesize an alternative, N-terminally truncated, polypeptide. Such primary and alternative proteins may have related but specialized functions. One of the first examples of internal initiation have been found within the *Escherichia coli* gene *infB* encoding translation initiation factor 2 (IF2). Initiation of *infB* translation either at the pTIS or iTIS leads to production of two IF2 isoforms with potentially distinct roles in cell physiology (Miller and Wahba, 1973; Nyengaard et al., 1991; Sacerdot et al., 1992). The products of in-frame internal initiation at several other bacterial genes have been reported to participate in various cellular functions ranging from virulence, to photosynthesis, or antibiotic production among others (reviewed in (Meydan et al., 2018)). In very few known cases, iTIS directs translation in a reading frame different from the primary ORF (D’Souza et al., 1994; Feltens et al., 2003; Yuan et al., 2018). The amino acid sequence and likely the function of a protein whose translation starts from an out-of-frame (OOF) internal start codon of a gene, would completely deviate from those of the primary protein (Feltens et al., 2003).

The majority of the few reported examples of translation of a protein from an iTIS have been discovered accidentally. Doubtlessly, most of the cases of alternative translation initiation remain enigmatic primarily due to the lack of systematic and robust approaches for detecting true start codons in bacterial genomes. Although several computational algorithms have successfully predicted the pTIS of many bacterial ORFs (Gao et al., 2010; Giess et al., 2017; Makita et al., 2007; Salzberg et al., 1998), iTIS prediction remains by far more challenging and has not even been pursued in most of those studies. The recent advent of new mass-spectrometry-based approaches have allowed the identification of N-terminal peptides of a range of proteins expressed in some bacterial species (Bienvenut et al., 2015; Impens et al., 2017). The resulting datasets revealed some peptides that likely originated from proteins whose translation initiates at an iTIS. However, the utility of the available mass-spectrometry techniques is intrinsically restricted due to the stringent requirements for the chemical properties, size, and abundance of the peptides that can be detected (Berry et al., 2016). Therefore, these methodologies had so far only limited success in identifying the protein products whose translation starts at an iTIS.

Ribosome profiling (Ribo-seq), based on deep sequencing of ribosome protected mRNA fragments (“ribosome footprints”), allows for genome-wide survey of translation (Ingolia et al., 2009). Ribo-seq experiments carried out with eukaryotic cells pre-treated with the translation initiation inhibitors harringtonine (Ingolia et al., 2011) and lactimidomycin (Gao et al., 2015; Lee et al., 2012) or with puromycin (Fritsch et al., 2012) showed specific enrichment of ribosome footprints at start codons of ORFs and facilitated mapping of TISs in eukaryotic genomes. These studies also revealed active translation of previously unknown short ORFs in the 5’ UTRs of many genes, significantly expanding the repertoire of the regulatory upstream ORFs. In addition, the presence of TISs within some of the coding regions in eukaryotic genomes was noted, where they were attributed primarily to leaky scanning through the primary start codons (Lee et al., 2012). Analogous studies, however, have been difficult to conduct in bacteria because of the paucity of inhibitors with the required mechanism of action. An inhibitor useful for mapping start sites should allow the assembly of the 70S translation initiation complex at a TIS but must prevent the ribosome from leaving the start codon. Unfortunately, most of the ribosomal antibiotics traditionally viewed as initiation inhibitors do not satisfy these criteria. Some of them (e.g. edeine or kasugamycin), interfere with binding of the ribosome to mRNA (Wilson, 2009) and thus are of little utility for TIS mapping. Other antibiotics, like thiostrepton or hygromycin A, may inhibit initiation of protein synthesis (Brandi et al., 2004; Polikanov et al., 2015), but these effects are only collateral to their ability of arresting the elongating ribosome (Cameron et al., 2002; Guerrero and Modolell, 1980) and, therefore, these compounds cannot be used for selective mapping of the sites of translation initiation on mRNAs. Recently, tetracycline (TET), an antibiotic that prevents aminoacyl-tRNAs from entering the ribosomal A site (Blanchard et al., 2004; Brodersen et al., 2000; Pioletti et al., 2001) and commonly known as an elongation inhibitor (Cundliffe, 1981), was used in conjunction with Ribo-seq to globally map pTISs in the *E. coli* genome (Nakahigashi et al., 2016). Although TET Ribo-seq data successfully revealed the pTISs of many of the actively translated genes, identification of iTISs was not performed in that work. Moreover, because TET can inhibit not only initiation, but also elongation of translation (Cundliffe, 1981; Orelle et al., 2013), it is impossible to distinguish whether the TET-generated peaks of ribosome density within the ORFs represented paused elongating ribosomes or ribosome initiating at an iTIS (Nakahigashi et al., 2016). Therefore, identifying an inhibitor that can specifically and selectively arrest the initiating ribosomes would provide an invaluable tool for mapping all TISs, primary or internal, in bacterial genomes.

Here we show that the antibiotic retapamulin (RET), a representative of the pleuromutilin class of protein synthesis inhibitors, exclusively stalls the ribosomes at the start codons of the ORFs. Brief pre-treatment of *E. coli* cells with RET dramatically rearranges the distribution of ribosomes along the ORFs, confining the ribosomal footprints obtained by Ribo-seq to the TISs of the genes. Strikingly, the application of the Ribo-seq/RET (Ribo-RET) approach to the analysis of bacterial translation revealed that more than a hundred of *E. coli* genes contain actively used iTISs. In vitro and in vivo experiments confirmed initiation of translation at some of the discovered iTISs and show that internal initiation may lead to production of proteins with distinct functions. Evolutionary conservation of some of the iTISs provide additional support for their functional significance. The obtained data show that alternative initiation of translation is widespread in bacteria and reveal a possible existence of the previously cryptic fraction of the bacterial proteome.

## RESULTS

### RET arrests the initiating ribosome at the start codons

Pleuromutilin antibiotics, including clinically-used semi-synthetic RET, bind in the peptidyl transferase center (PTC) of the bacterial ribosome where they hinder the placement of the P- and A-site amino acids (Figure S1A and S1B), thus preventing peptide bond formation (Davidovich et al., 2007; Poulsen et al., 2001; Schlunzen et al., 2004). In vitro studies have shown that presence of fMet-tRNA and RET in the ribosome are not mutually exclusive (Yan et al., 2006). Therefore, we reasoned that RET may allow the assembly of the 70S initiation complex at the start codon, but by displacing the aminoacyl moiety of the initiator fMet-tRNA and interfering with the placement of the elongator aminoacyl-tRNA in the A site, it could prevent formation of the first peptide bond.

The results of polysome analysis was compatible with the view of RET being a selective inhibitor of translation initiation, because treatment of *E. coli* cells with high concentrations of RET, 100-fold over the minimal inhibitory concentration (MIC), led to a rapid conversion of polysomes into monosomes (Figure S1C). We then carried out toeprinting experiments (Hartz et al., 1988; Orelle et al., 2013) in order to test whether RET could indeed capture ribosomes at start codons. When model genes were translated in an *E. coli* cell-free system (Shimizu et al., 2001), addition of RET resulted in ribosome stalling exclusively at the start codons of the ORFs (Figure 1A, ‘RET’ lanes), demonstrating that this antibiotic readily, and possibly specifically, inhibits translation initiation. In contrast, TET, which was used previously to map TISs in the *E. coli* genome (Nakahigashi et al., 2016), halted translation not only at the translation initiation sites but also at downstream codons of the ORFs (Figure 1A, ‘TET’ lanes), confirming its ability to interfere with both initiation and elongation phases of protein synthesis (Orelle et al., 2013).

**Figure 1:**
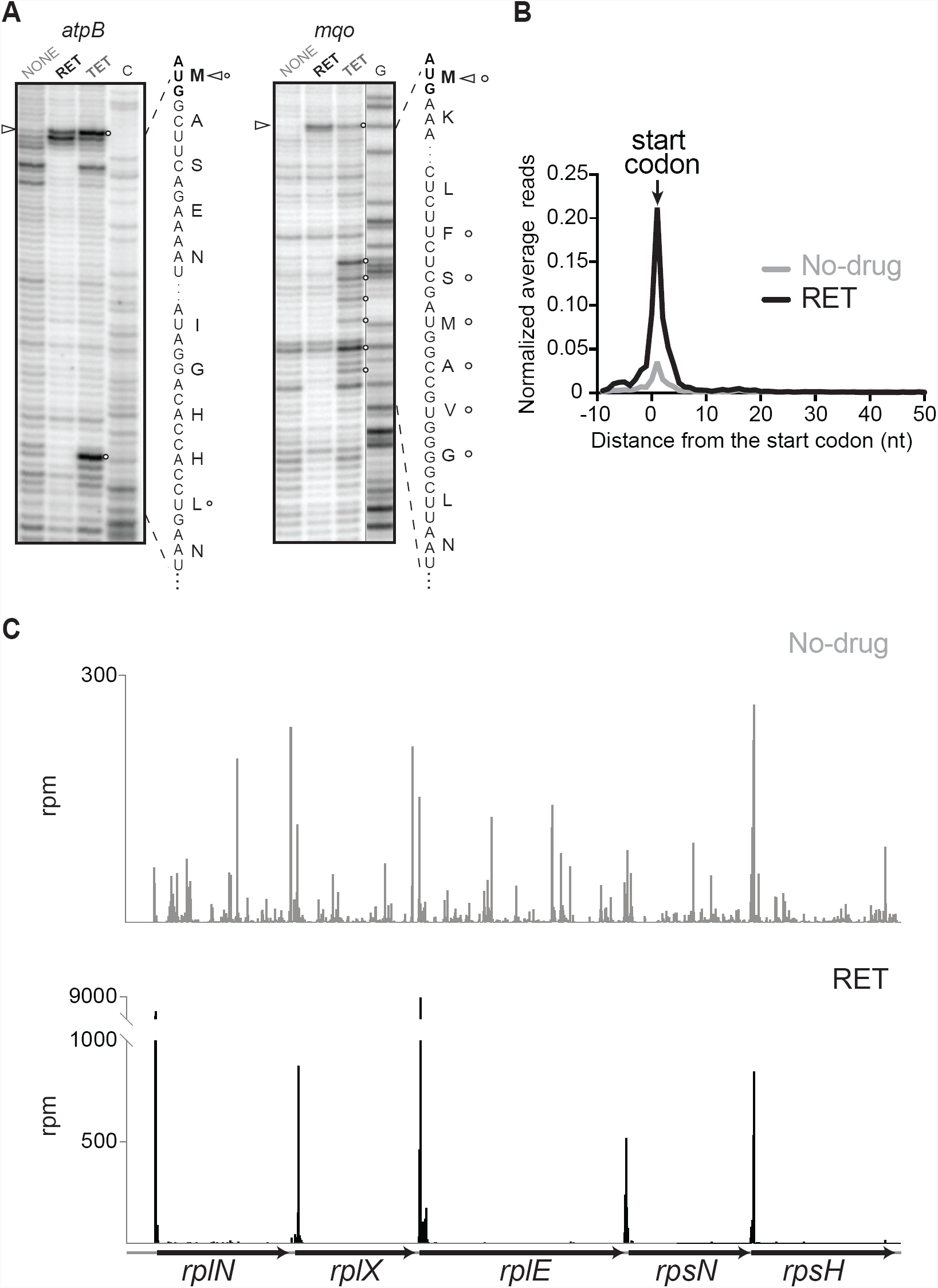
RET specifically arrests ribosomes at translation initiation sites. (A) Toeprinting analysis of RET- and TET-mediated ribosome stalling during cell-free translation of two model *E. coli* genes. The circles indicate TET-induced translation arrest sites, whereas triangles designate RET-mediated translational stalls. The lane labeled NONE represents samples translated in the absence of antibiotics. Sequencing lanes (C for the *atpB* gene and G for the *mqo* gene) were used to map the sites of drug-induced arrests. The nucleotide sequences of the genes and the encoded proteins are shown. Note that the toeprint bands correspond to the mRNA residue 16-17 nt downstream from the first nucleotide of the codon residing in the P site of the arrested ribosome. (B) Metagene analysis plot representing normalized average relative density reads in the vicinity of the annotated start codons of the *E. coli* genes in exponentially growing cells treated with RET (black line) or not exposed to antibiotic (gray line). (C) Redistribution of ribosome footprints density within the *spc* operon upon RET treatment. Top profile: no-drug sample; bottom profile: RET-treated cells. See also Figures S1.

The outcomes of the polysome- and toeprinting analyses, and structural data showing the incompatibility of the extended nascent protein chain with RET binding (Figure S1B), encouraged us to assess by Ribo-seq whether RET can be used for mapping translation start sites in the living bacterial cells. Even though a brief 2 min exposure of the Δ*tolC* derivative of the *E. coli* strain BW25113 to a 32-fold MIC of RET was sufficient to nearly completely halt protein synthesis (Figure S1D), Ribo-seq experiments were carried out by incubating cells with 100-fold MIC concentration of the antibiotic for 5 min in order to ensure efficient drug binding and providing the elongating ribosomes enough time to complete translation of even long or slowly-translated mRNAs prior to cell harvesting. Analysis of the resulting Ribo-seq data showed that the treatment of *E. coli* with RET led to a striking redistribution of the translating ribosomes along the ORFs. The occupancy of the internal and termination codons of the expressed genes was severely reduced compared to the untreated control, whereas the ribosome density peaks at the start codons dramatically increased (Figure 1B and 1C). Although a generally similar trend can be observed in the metagene analysis of the Ribo-seq data in the RET-(this paper) and TET-treated cells (Nakahigashi et al., 2016), the start-codon peak in the TET experiments is smaller and broader compared to the peak of the RET-stalled ribosomes (Figure S1E and S1F), reflecting a higher potency of RET as initiation inhibitor. By applying fairly conservative criteria (see STAR Methods), we were able to detect the distinct peaks of ribosome density at the annotated start codons (pTISs) of 991 out of 1153 (86%) *E. coli* genes expressed in the BW25113 strain under our experimental conditions in the absence of antibiotic. The 86% success rate in pTIS mapping confirmed the power of the approach for identifying translation start sites in bacterial genes. The magnitude of the start codon peaks at the remaining 14% of the genes did not pass our strict ‘minimal number of reads’ threshold (see STAR Methods) likely reflecting in part possible changes in gene expression upon addition of RET. The number of the recognizable pTISs could be evidently expanded by increasing the depth of sequencing of the Ribo-seq libraries.

Taken together, our in vitro and in vivo results showed that RET acts as a specific inhibitor of bacterial translation initiation and in combination with Ribo-seq can be used for mapping the pTISs of the majority of actively translated genes in bacterial genomes. We named the Ribo-seq/RET approach ‘Ribo-RET’.

### Ribo-RET unmasks initiation of translation at internal codons of many bacterial genes

The majority of the ribosome footprints in the Ribo-RET dataset mapped to the annotated pTISs. Strikingly, in a number of genes we also observed peaks at certain internal codons (Figure 2A). Hypothetically, the presence of internal Ribo-RET peaks could be explained by pausing of elongating ribosomes at specific sites within the ORF, if the stall does not resolve during the time of antibiotic treatment. Nonetheless, this possibility seemed unlikely, because we did not detect any substantial Ribo-RET peak even at the most prominent programmed translation arrest site in the *E. coli* genome within the *secM* ORF (Nakatogawa and Ito, 2002) (Figure S1G). The likelihood that the internal RET peaks originated from context-specific translation elongation arrest observed with some other protein synthesis inhibitors (Kannan et al., 2014; Marks et al., 2016) was similarly implausible because biochemical (Dornhelm and Hogenauer, 1978) and structural (Davidovich et al., 2007) data strongly argue that RET cannot bind to the elongating ribosome (Figure S1B). We concluded, therefore, that the Ribo-RET peaks at internal sites within ORFs must represent ribosomes caught in the act of initiating translation.

**Figure 2:**
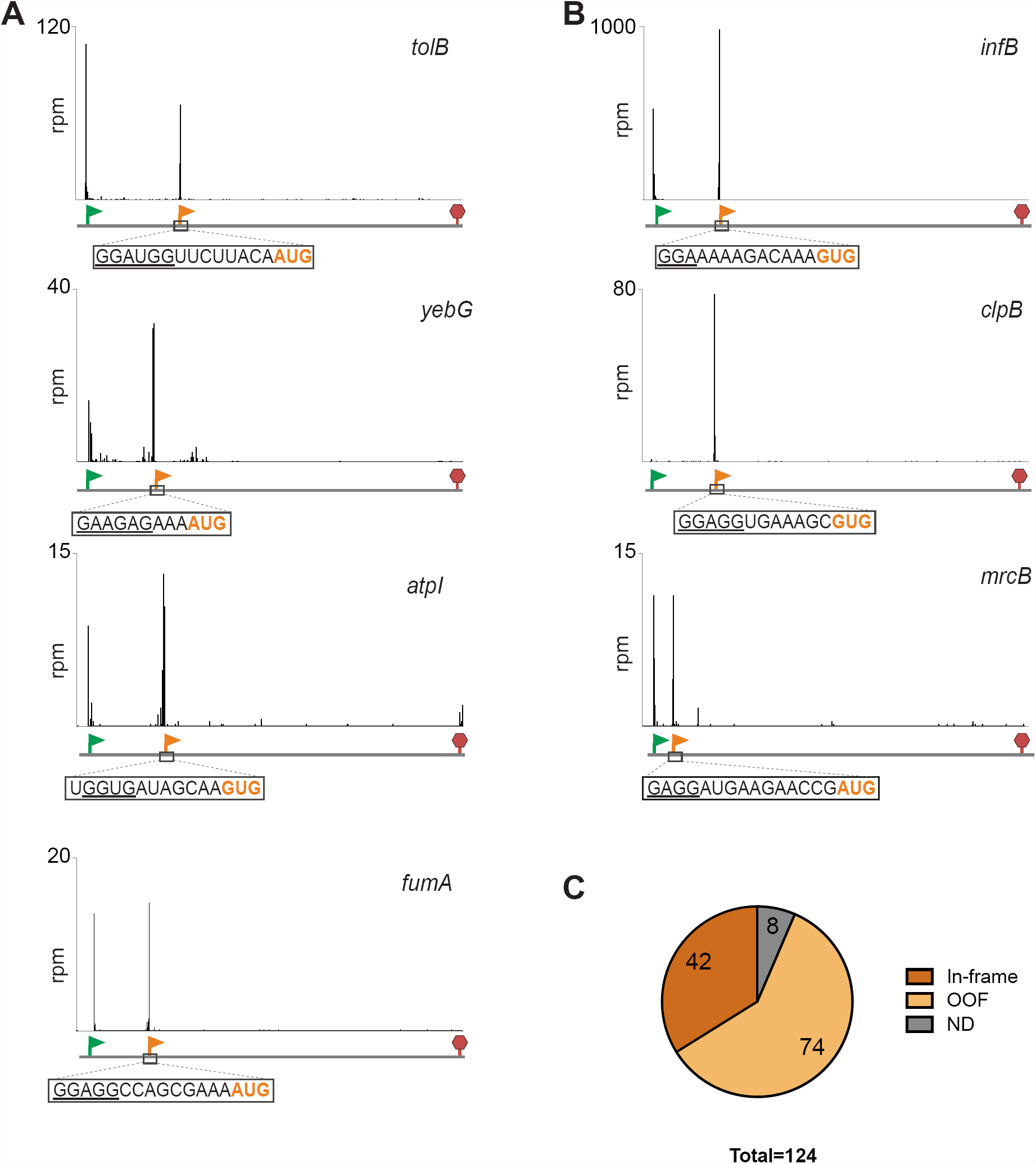
Ribo-RET reveals the presence of iTISs in many bacterial genes. (A) Examples of Ribo-RET profiles demonstrating the presence of well-pronounced ribosome density peaks not only at the annotated pTISs (marked with green flags) but also at internal codons of *E. coli* genes (orange flags) not known previously to contain iTISs. The putative start codon of the iTIS is highlighted in orange and the SD-like sequence is underlined. (B) Ribo-RET profiles of the three *E. coli* genes, *infB, clpB, mrcB*, where iTISs had been previously characterized. (C) The iTISs common between the *E. coli* K-strain BW25113 and the B-strain BL21. The sector labeled as ND (not determined) represents the Ribo-RET peaks observed at internal ORF sites without any previously recognized start codon in the vicinity. See also Table S1.

Three *E. coli* genes, *infB, mcrB and clpB,* have been previously reported to encode two different polypeptide isoforms; in each of these genes the translation of the full-size protein is initiated at the pTIS while the shorter isoform is expressed from an iTIS (Broome-Smith et al., 1985; Park et al., 1993; Plumbridge et al., 1985). Examination of the Ribo-RET profile within these genes showed well-defined and highly-specific peaks of ribosome density (Figure 2B) at the previously experimentally verified iTISs. This result proved the utility of Ribo-RET for mapping iTISs in bacterial genes and provided independent evidence that the RET peaks at internal codons represent initiating ribosomes.

Among the *E. coli* BW25113 genes expressed in our conditions, we identified 239 that contain at least one iTIS characterized by the presence of a ribosome density peak whose size is at least 10% relative to the pTIS peak in the same gene. To further expand the systematic identification of *E. coli* genes with internal translation start sites, we applied the Ribo-RET approach to the Δ*tolC* derivative of the BL21*E. coli* strain, the B-type variant which is genetically distinct from the K-strain BW25113 (Grenier et al., 2014; Studier et al., 2009) used in the original Ribo-RET experiment. Similar to our findings with the BW25113 cells, Ribo-RET analysis identified a number of genes in the BL21 strain with well-defined iTISs. Of these, 124 iTISs were common between the two strains (Table S1). Additional 115 Ribo-RET iTIS peaks exclusively identified in the BW25113 cells and 496 Ribo-RET peaks specific for the BL21 cells may represent strain-specific iTISs candidates (Table S1). We limited our subsequent analysis to the iTISs conserved between the two strains, among which, 42 directed translation in frame with the main gene whereas start codons of the remaining 74 iTISs were out of frame relative to the main ORF (Figure 2C and Table S1). In the following section we will consider these two classes separately.

### Internal translation initiation sites that are in frame with the main ORF

The in-frame iTISs exploit various initiator codons that have been shown previously to be capable of directing translation initiation in *E. coli* (Chengguang et al., 2017; Hecht et al., 2017), although similar to the pTISs, the AUG codon is the most prevalent (Figure 3A). A SD-like sequence could be recognized in front of many of in-frame internal start codons (Table S1).

**Figure 3:**
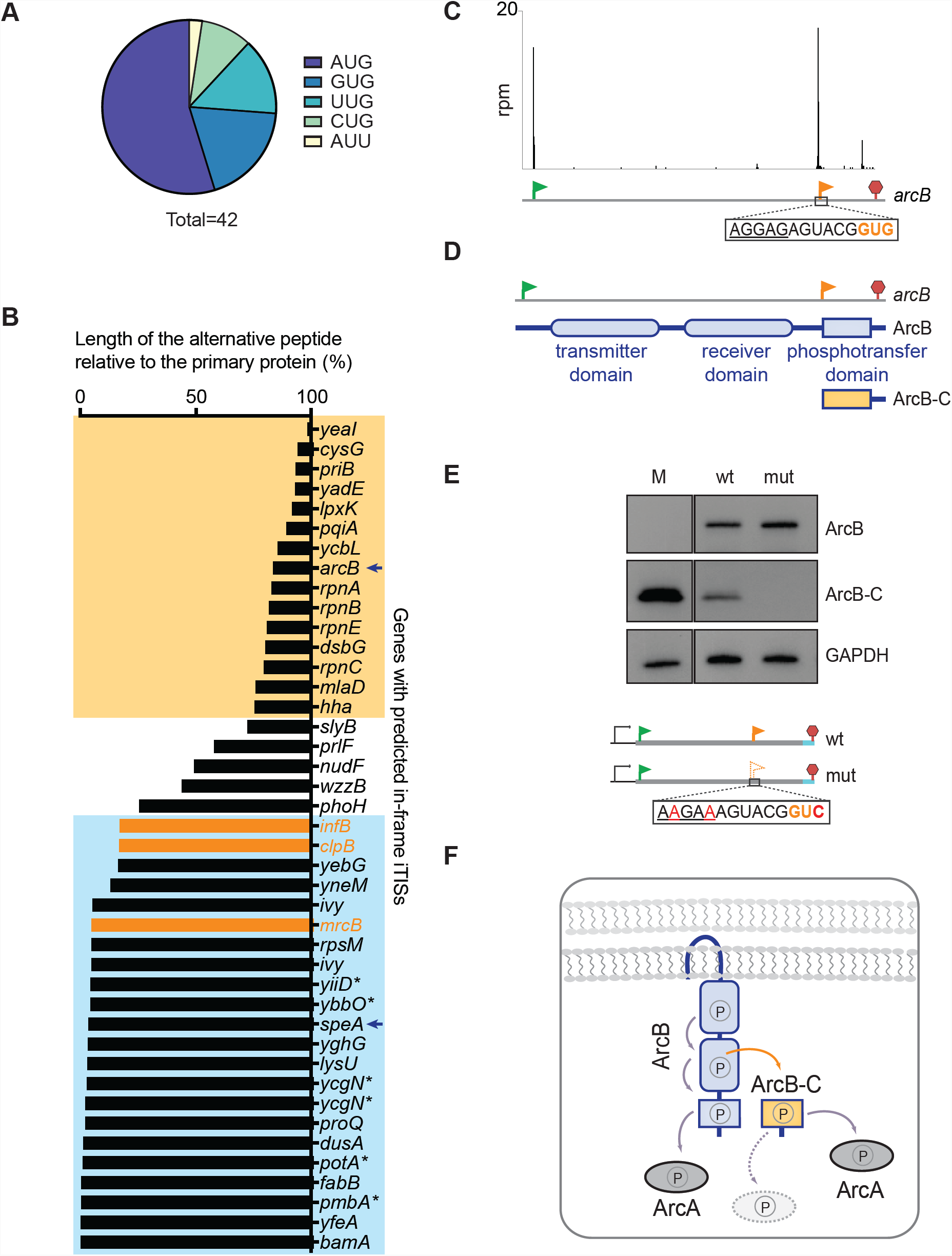
In-frame internal initiation can generate N-terminally truncated products that could play physiological roles. (A) The relative use of various putative start codons at the identified in-frame iTISs. (B) The length of the predicted alternative proteins whose translation is initiated at the in-frame iTISs relative to the corresponding primary proteins. The previously known examples of genes with in-frame iTISs (shown in Figure 2B) are shown in orange. The genes with the iTISs located within the 3’ or 5’ quartile of the gene length are highlighted with yellow or blue boxes, respectively. The candidate genes where, based on TET-assisted Ribo-seq, re-annotation of their pTISs was proposed (Nakahigashi et al., 2016) are indicated by asterisks. The *arcB* and *speA* genes discussed in more detail are indicated by blue arrows. (C) The Ribo-RET profile of the *arcB* gene shows a ribosomal density peak at the pTIS (green flag) and at the iTIS candidate (orange flag). The iTIS start codon is highlighted in orange and the SD-like sequence is underlined. (D) Schematic representation of the full-length ArcB protein bearing its three functional domains. The putative protein product, ArcB-C, resulting from initiation of translation at the *arcB* iTIS revealed by Ribo-RET, would specifically encompass the phosphotransfer domain. (E) Western blot analysis of the C-terminally 3XFLAG-tagged translation products of the *arcB* gene expressed from a plasmid in *E. coli* cells. The ArcB and ArcB-C proteins are expressed from the wild-type *arcB* (lane wt); inactivation of iTIS by the indicated mutations abrogate production of ArcB-C (lane mut). The marker protein representing the 3XFLAG ArcB-C segment of ArcB (lane M) was expressed from a specially-constructed plasmid (see STAR Methods). (F) Possible role of the ArcB-C protein translated from the *arcB* iTIS. The phosphorelay across the domains of membrane-embedded ArcB (gray arrows) results in the activation of the response regulator ArcA, which then triggers the expression of genes critical for survival under low-oxygen conditions (Alvarez et al., 2016). The product of internal initiation, ArcB-C, could improve the signal amplification capabilities of the ArcBA system and, being cytosolic and detached from the membrane-bound ArcB, could activate additional response regulators (represented by a dotted oval). See also Figure S2.

A polypeptide whose synthesis is initiated at an in-frame iTIS would represent an N-terminally truncated form of the primary protein. The candidate proteins resulting from in-frame internal initiation encompass a wide range of sizes, from 6 to 805 amino acids in length (Table S1). The locations of the in frame iTISs within the main ORFs vary but, in general, they show a bimodal distribution with a large number of internal start sites clustering close to the beginning of the gene while others are within the 3’ terminal quartile of the ORF length (Figure 3B). As a representative example of a 3’-proximal iTIS, we selected that of the *arcB* gene, while the iTIS of *speA* was examined as the case of a 5’-proximal start site.

### 3’-proximal internal initiation can generate an isoform of the primary protein with specialized functions

The gene *arcB* encodes the membrane-anchored sensor kinase ArcB of the two-component signal transduction system ArcB/A that helps bacteria to sense and respond to changes in oxygen concentration (Alvarez and Georgellis, 2010) (Figure 3C and 3D). The ArcB protein consists of the transmitter, receiver and phosphotransfer domains (Figure 3D). Under microaerobic conditions, ArcB undergoes a series of phosphorylation steps and eventually transfers a phosphoryl group to the response regulator ArcA that controls expression of nearly 200 genes (Alvarez and Georgellis, 2010; Salmon et al., 2005). ArcB-C, the C-terminal phosphotransfer domain of the membrane-bound ArcB is the ultimate receiver of the phosphoryl group within the ArcB protein and serves as the phosphoryl donor for ArcA (Alvarez et al., 2016).

Our Ribo-RET data showed a strong ribosome density peak at an iTIS in *arcB*, with the putative start codon GUG located precisely at the boundary of the segment encoding the ArcB-C domain (Figure 3C, D). This observation suggested that initiation of translation at the *arcB* iTIS could generate a diffusible ArcB-C polypeptide corresponding to the C-terminal domain of the membrane-anchored ArcB kinase (Figure 3D). To test this possibility, we introduced the 3xFLAG-coding sequence at the 3’ end of the *arcB* gene, expressed it from a plasmid in *E. coli* cells and analyzed the protein products by Western-blotting. Expression of the tagged *arcB* resulted in the simultaneous production of two proteins: the full-size ArcB and a second, smaller polypeptide, whose apparent molecular weight of 14 kDa is consistent with that of FLAG-tagged ArcB-C (Figures 3E and S2A). Disruption of the *arcB* iTIS by synonymous mutations at the start codon and the upstream SD-like sequence did not affect the synthesis of the full-length ArcB but abrogated the production of ArcB-C (Figure 3E). These results demonstrate that the iTIS within the *arcB* gene may indeed be utilized in the cell for synthesis of a stand-alone ArcB-C protein, detached from the membrane-bound ArcB (Figure 3F). It has been previously shown that the isolated ArcB-C transmitter domain is capable of serving in vitro as a phosphoryl acceptor for the ArcB-catalyzed phosphorylation reaction and then can catalyze the phosphoryl transfer to ArcA (Alvarez and Georgellis, 2010), supporting the idea that a self-standing ArcB-C protein is likely also functional in vivo. Diffusible ArcB-C may either facilitate the operation of the ArcB-ArcA signal transduction pathway or could possibly phosphorylate other response regulators enabling a cross-talk with other signal transduction systems of the cell (Yaku et al., 1997) (Figure 3F). The functional significance of producing the full-size ArcB sensory kinase and an independent ArcB-C phosphotransfer domain from the same gene is further supported by preservation of the *arcB* iTIS among several bacterial species (Figure S2B).

The expression of ArcB-C from the *arcB* iTIS is apparently quite efficient because in *E. coli* and *Salmonella enterica* Ribo-seq datasets obtained in the absence of RET, an upshift in the ribosome density can be observed at the *arcB* codons located downstream from the iTIS (Baek et al., 2017; Kannan et al., 2014; Li et al., 2014) (Figure S2C-S2F). Curiously, the average ribosome occupancy of the *arcB* codons before and after the iTIS vary under different physiological conditions (Figure S2E and S2F), suggesting that the relative efficiency of utilization of the *arcB* pTIS and iTIS could be regulated.

Another remarkable example of a 3’-proximal iTIS that likely directs production of a functional protein is found in homologous *rpnA, rpnB, rpnC, rpnD* and *rpnE* genes found in the *E. coli* genome that encode nucleases involved in DNA recombination (Kingston et al., 2017). All the *rpn* genes in the *E. coli* show Ribo-RET peaks corresponding to in-frame iTISs that appear to be the major initiation sites of each of these genes (Figure S2G), at least under the growth conditions of our experiments. Strikingly, an alternative polypeptide expressed from an in-frame iTIS in one of these genes (*rpnE*) is 98% identical to the product of an independent *ypaA* gene (Figure S2H) revealing distinct functionality of the products of the internal initiation within the genes of the *rpn* family.

### 5’-proximal iTIS gene may generate differentially-targeted proteins

Twenty-two (52%) of the conserved in-frame *E. coli* iTISs are within the first quarter of the length of the main gene. In 18 of these 22 genes, the iTIS is located within the first 10% of the ORF length (Figure 3B). Compared to the main protein product of a gene, an alternative polypeptide generated from a 5’-proximal iTIS would lack only few N-terminal amino acid residues. The functions of such a protein would likely be very similar to those of the primary gene product, raising the question of whether the presence of a secondary, 5’-proximal, iTIS within the gene could benefit the cell.

We examined the Ribo-RET-detected 5’-proximal iTIS of *speA*, the gene that encodes arginine decarboxylase, an enzyme involved in polyamine production (Michael, 2016). Arginine decarboxylase (SpeA) has been found in the *E. coli* cytoplasmic and periplasmic fractions (Buch and Boyle, 1985) and was reported to be represented by two polypeptide isoforms, SpeA-74, with an approximal molecular weight of 74 kDa, and a smaller one of ∼ 70 kDa, SpeA-70 (Buch and Boyle, 1985; Wu and Morris, 1973). The latter protein was suggested to be a product of co-secretional maturation of the full-length SpeA-74 (Buch and Boyle, 1985). Our analysis, however, revealed two Ribo-RET peaks in the *speA* ORF: one corresponding to the annotated pTIS and the second one mapped to an iTIS at codon Met-26 (Figure S3A). Initiation of translation at the pTIS and iTIS of *speA* would generate the 73,767 Da and 71,062 Da forms of SpeA, respectively, arguing that the SpeA-70 isoform is generated due to initiation of translation at the *speA* iTIS. In support of this conclusion, the peptide (M)SSQEASKMLR, which precisely corresponds to the N-terminus of the short SpeA isoform defined by Ribo-RET, can be found in the database of the experimentally-identified *E. coli* N-terminal peptides (Bienvenut et al., 2015).

Previous studies suggested that SpeA-74 is targeted to the periplasm due to the presence of a putative N-terminal secretion signal sequence (Buch and Boyle, 1985). A segment of this signal sequence would be missing in the SpeA-70 isoform and therefore this shorter polypeptide would be confined to the cytoplasm (Figure S3B). Therefore, utilization of the 5’-proximal iTIS of *speA* would result in retaining a fraction of SpeA enzyme in the cytoplasm. The 5’-proximal iTISs identified in some other *E. coli* genes encoding secreted proteins (e.g. *bamA, ivy* or *yghG* (Stegmeier and Andersen, 2006; Strozen et al., 2012; Wasinger and Humphery-Smith, 1998), may serve similar purposes since initiation from the internal start codons would partially eliminate the predicted signal sequences from the polypeptide (Figure S3C). A similar strategy for targeting polypeptide isoforms to different cellular compartments has been described for few other bacterial proteins (reviewed in (Meydan et al., 2018)).

Six of the 5’-proximal iTISs (marked by asterisks in Figure 3B) have been detected previously by TET Ribo-seq and suggested to represent incorrectly annotated primary translation start sites (Nakahigashi et al., 2016). Some of the 5’-proximal internal initiation site candidates may indeed correspond to mis-annotated pTISs because no ribosome density peaks were observed at the pTISs of the corresponding genes in our Ribo-RET datasets (Table S1). However, such conclusion could be drawn only cautiously because the utilization of the upstream pTIS could depend on the growth conditions.

The comparison of Ribo-RET-identified in-frame iTISs among two *E. coli* strains show that many of them are strain specific, indicating high variability of the internal translation initiation landscape. Nevertheless, the evolutionary pressure is expected to impose some degree of preservation upon the iTISs that provide a competitive advantage for the cell. One such example is preservation of the iTIS within the *arcB* gene discussed earlier.

To assess conservation of other iTISs, we analyzed alignments of bacterial genes homologous to the *E. coli* genes containing internal in-frame start sites. Sequence logos and codon conservation plots indicated preservation of in-frame potential initiation sites and locally enhanced synonymous site conservation for several of the genes, including *phoH, speA, yebG, yfaD* and *yadD* (Figure S4A-S4E). However, it remains to be seen whether these conserved regions are relevant to promoting iTIS usage or simply represent unrelated cis-elements embedded within these genes. The other iTISs identified by Ribo-RET in the *E. coli* genome show a lower degree of evolutionary conservation and could have arisen in response to species-specific needs.

### Ribo-RET identified iTISs that are out of frame relative to the main ORF

Only two examples of OOF internal initiation in bacterial genes had been previously described: the *comS* gene in *Bacillus subtilis*, that is nested within the *srfAB* gene (D’Souza et al., 1994; Hamoen et al., 1995), and the *rpmH* gene, which in *Thermus thermophilus* resides within the *rnpA* gene (Feltens et al., 2003) (reviewed in (Meydan et al., 2018)). Our Ribo-RET analysis showed that 74 OOF iTISs are potentially exploited as alternative translation start sites in the examined *E. coli* BW25113 and BL21 strains. The majority of the predicted OOF iTISs contain AUG as a start codon, but other known initiation codons are also found with the frequencies resembling those in pTISs or in in-frame iTISs (Figure S5A). The location of the OOF iTISs varies significantly between the host genes and the peptides generated by translation initiated at the predicted OOF iTISs would range in size from 2 to 84 amino acids (Figures 4A and S5B).

**Figure 4:**
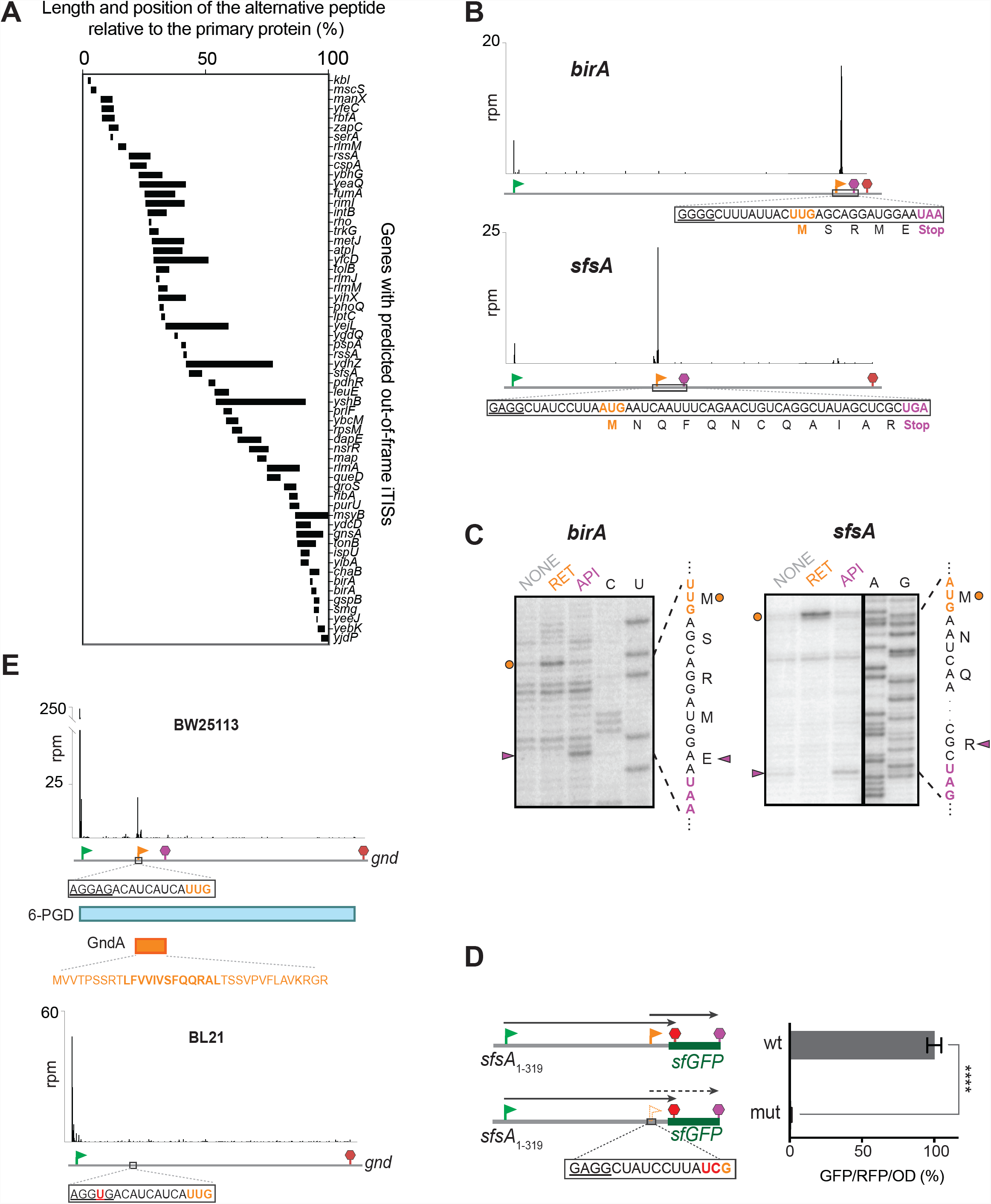
OOF iTISs revealed by Ribo-RET can direct initiation of translation. (A) The length and location of the predicted OOF alternative protein-coding segments relative to the main ORF. (B) Ribo-RET profiles of *birA* and *sfsA* as representative genes with putative OOF iTISs. The pTIS and OOF iTIS start codons are shown by green and orange flags, respectively. The main frame and alternative frame stop codons are indicated by red and purple stop signs, respectively. The entire sequence of the alternative ORF is shown with the start codon highlighted in orange, stop codon highlighted in purple and putative SD-sequence underlined. (C) Toeprinting analysis revealing antibiotic-induced stalling of the ribosome at the start and stop codons of the alternative ORFs within the *birA* and *sfsA* genes. RET (lanes marked RET) arrests ribosomes at the start codons of the alternative ORFs, whereas translation termination inhibitor Api137 induces translation arrest at the stop codons of those ORFs (lanes marked API). No antibiotic was present in the samples ran in lanes marked NONE. The templates used in toeprinting analysis contained the sequences of the full-length *birA* or *sfsA* genes. The nucleotide and amino acid sequences of the alternative ORFs are shown. Internal OOF start and stop codons are indicated by orange circles and purple triangles, respectively. Sequencing lanes are indicated. (D) The OOF iTIS within *sfsA* initiates translation in the cell. Schematic representation of the plasmid-based reporter construct (the reference *rfp* gene is not shown). The GFP-coding sequence is OOF relative to the pTIS start codon (green flag) and its expression is controlled by the iTIS (orange flag). Mutation of the iTIS AUG start codon to UCG alleviates the expression of the reporter. The first stop codon in-frame with the pTIS is shown by the red stop sign and the stop codon in-frame with the iTIS is indicated by the purple stop sign. The bar graph shows the change in relative green fluorescence in response to the iTIS start codon mutation. (E) Ribo-RET snapshots of the *gnd* gene in the *E. coli* BW25113 strain (top) that reveals the location of the start codon of the alternative ORF (marked by the orange flag for the start codon and purple stop sign for the stop codon). The amino acid sequence of the alternative ORF product GndA is shown in orange. The sequence of the GndA tryptic peptide identified by mass spectrometry (Yuan et al., 2018), is indicated with bold characters. The snapshot of the *gnd* gene of the BL21 strain shows the lack of the RET-generated density peak at the iTIS site, presumably due to a point mutation (red, underlined) in the SD-like region of the iTIS. See also Figure S5 and Table S1.

We selected two of the Ribo-RET-identified OOF iTIS candidates to examine if they indeed can direct initiation of translation. The *E. coli* genes *sfsA* and *birA* encode transcription regulators; *birA* also possesses biotin ligase activity (Eisenberg et al., 1982; Kawamukai et al., 1991). In the *birA* gene, the presence of an OOF UUG internal start site (overlapping the Leu_300_ codon of the main frame) would direct translation of a 5-amino acid long peptide (Figure 4B). Internal initiation at the OOF AUG of the *sfsA* gene (overlapping the Leu_95_ codon of the main ORF) would generate a 12 amino acid long peptide (Figure 4B). When the full-size *sfsA* and *birA* genes were translated in vitro, addition of RET to the reaction resulted not only in the appearance of toeprint bands at the pTISs of the corresponding genes (Figure S5C and S5D), but also at the OOF iTISs identified by Ribo-RET (Figures 4C, lanes ‘RET’, orange dots). Furthermore, the addition of the translation termination inhibitor Api137 (Florin et al., 2017) to the translation reactions led to the appearance of toeprint bands at the stop codons corresponding to the OOF ORFs, indicating that the ribosomes not only bind to the OOF start codons but do translate the entire alternative ORF (Figure 4C, lanes ‘API’, magenta triangles).

We further examined whether the alternative ORF in the *sfsA* gene is translated in vivo. For this purpose, we designed a dual red/green fluorescent proteins (RFP/GFP) reporter plasmid, where translation of the *gfp* gene is initiated at the OOF iTIS within the *sfsA* gene (Figure 4D). Introduction of the reporter construct into *E. coli* cells resulted in active expression of the GFP protein (Figure 4D, right panel), indicating that the OOF iTIS within the *sfsA* gene is indeed utilized for initiation of translation. This conclusion was further corroborated when mutations in the internal AUG start codon abolished GFP production (Figure 4D). The verified functionality of the OOF iTIS in the *sfsA* gene suggests that either the encoded protein products or the mere act of translation of the OOF ORFs may play a functional role.

An independent validation of the functional significance of one of the OOF iTISs identified by Ribo-RET came from a recent study aimed at characterizing *E. coli* proteins activated by heat-shock (Yuan et al., 2018). Among the tryptic peptides, one mapped to the *gnd* gene, which encodes 6-phosphogluconate dehydrogenase (6-PGD). Remarkably, the sequence of the identified peptide was encoded in the -1 frame but the location of the start codon from which translation of the alternative protein (named GndA) would initiate remained ambiguous (Yuan et al., 2018). Our Ribo-RET data not only validated those findings, but also revealed that the GndA translation initiates most likely at the UUG codon (overlapping the Ile_97_ codon of *gnd*), which is preceded by a strong SD sequence (Figure 4E). Of note, the Ribo-RET peak that we observed at this iTIS was relatively small compared to the pTIS peak of the *gnd* gene; the efficiency of translation initiation at the iTIS likely becomes more pronounced under heat-shock conditions. Interestingly, the Ribo-RET signal that would reflect initiation of the *gndA* translation is absent in the Ribo-RET data collected with the BL21 strain, perhaps because due to a single nucleotide alteration within the SD sequence preceding the OOF iTIS (Figure 4E). This observation illustrates the strain-specificity of expression of functional alternative proteins. In line with this conclusion, most of the OOF iTISs identified in the *E. coli* genome do not exhibit any significant evolutionary conservation. The strongest example that exhibits near-threshold significance of the OOF iTIS conservation is the *tonB* gene that encodes the periplasmic component of the system involved in transport of iron-siderophore complex and vitamin B12. Furthermore, the internal initiation at this site is apparently sufficiently strong to be observed as an upshift of the local density of ribosome footprints even in the Ribo-seq data collected from *E. coli* cells not treated with antibiotic (Figure S4F).

### Start-Stop sites may modulate translation of the primary gene

Among the 74 OOF iTIS candidates, our Ribo-RET data revealed 14 unique sites where the start codon is immediately followed by a stop codon (Table S1). We called these unique sites “Start-Stops”. Although start-stops have been identified in the 5’ UTRs of some viral and plant genes and have been suggested to play regulatory functions (Krummheuer et al., 2007; Tanaka et al., 2016), operational Start-Stops have not been previously reported within the bacterial or archaeal genes.

We selected the identified start-stops within the genes *yecJ and hslR* (Figure 5A) for further analysis. Specifically, we tested whether the corresponding iTISs can indeed direct initiation of translation. In vitro studies, carried out using the full-length *yecJ* or *hslR* genes, showed that in both cases addition of initiation inhibitor RET or termination inhibitor Api137 caused the appearance of coincidental toeprint bands since, in these particular cases, either one of the inhibitors lead the ribosomes stalling at the initiation codon of the start-stop site (Figure 5B). These results demonstrated that the iTISs of the Start-Stops nested in the *yecJ* and *hslR* genes can direct ribosome binding. We then extended the analysis to living cells by fusing the *gfp* gene, devoid of its own start codon, immediately downstream from the AUG codon of the *yecJ* iTIS (but without its associated stop codon) (Figure 5C). When the resulting construct was expressed from a plasmid, GFP fluorescence was readily detectable as long as the initiator codon of the Start-Stop site was intact, but was significantly reduced when this AUG codon was mutated to ACG (Figure 5C). These results demonstrated that the start codon of the OOF start-stop site within the *yecJ* is operational in vivo.

**Figure 5:**
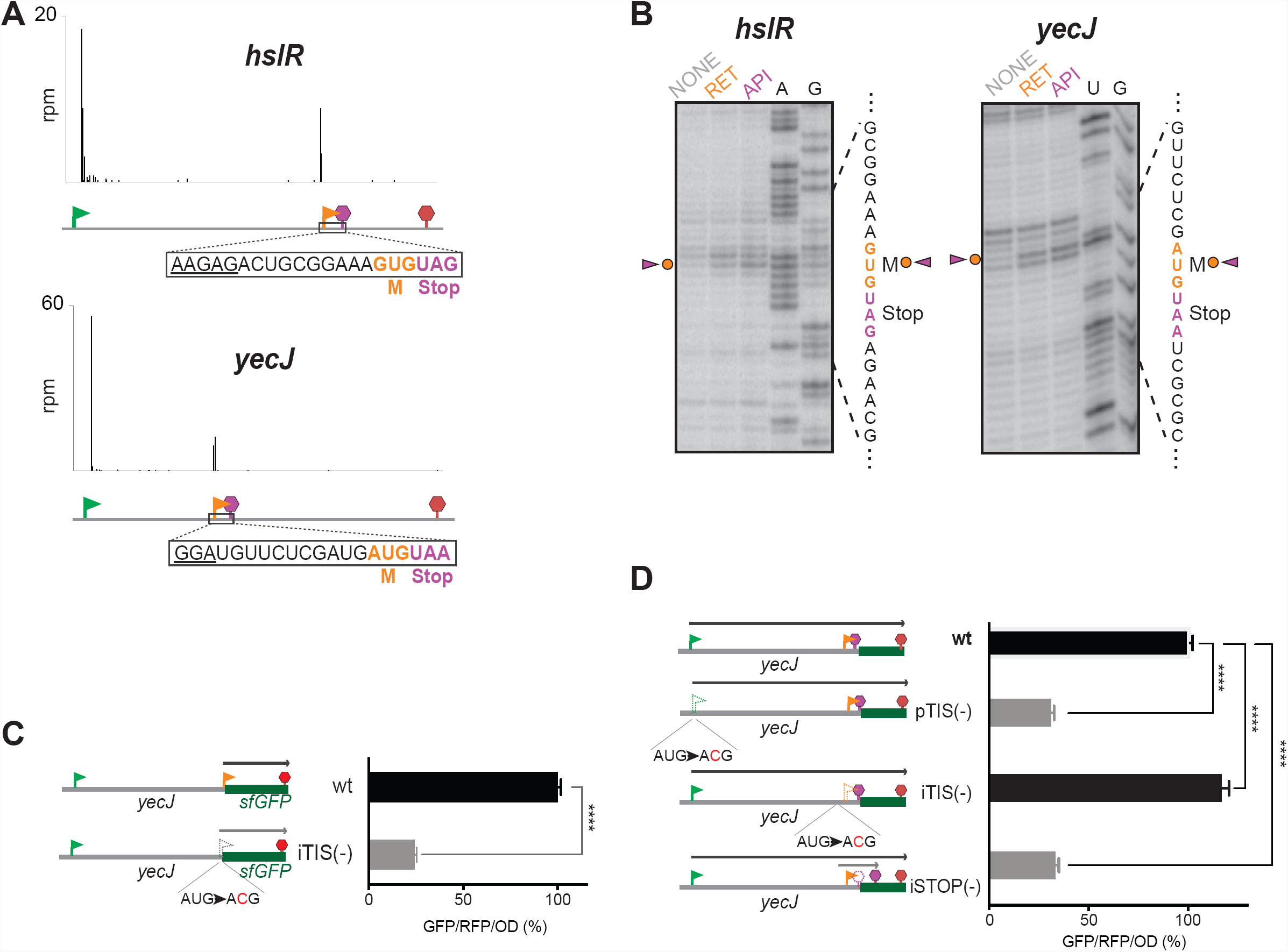
Start-Stops within *E. coli* genes. (A) Examples of *E. coli* genes with the Ribo-RET-revealed ribosome density at an OOF iTIS (orange flag) immediately followed by a stop codon (purple stop sign). The sequence of the minimal ORFs are shown, including the SD-like sequences preceding them. Start and stop codons of the primary ORF are indicated by green flags and red stop signs, respectively. (B) Toeprinting gels showing the ribosomes stalled at the start codons of the Start-Stop sites within the *hslR* and *yecJ* genes in response to the presence of translation initiation (RET) and translation termination (API) inhibitors. The nucleotide and amino acid sequences of the minimal ORFs are shown. The Start-Stop sites are indicated by orange circles and purple triangles. NONE indicates samples that contained no antibiotic. Sequencing lanes are indicated. (C) The start codon of the Start-Stop within the *yecJ* gene can direct initiation of translation in vivo. Left, schematic representation of the reporter constructs (the reference *rfp* gene is not shown) where GFP expression is directed by the start codon of the OFF iTIS (orange flag) of *yecJ*. Right, the bar graphs show the relative translation efficiency, as estimated by GFP/RFP/OD (%) ratio after 10 h induction of the reporter transcription. The expression of *gfp* is severely abrogated by a mutation that disrupts the start codon of the OOF iTIS (iTIS (-) construct). (D) Translation of the Start-Stop of *yecJ* impacts expression of the host gene. Left, schematic representation of the reporter constructs where the *yecJ* pTIS (green flag) controls expression of the YecJ-GFP chimeric protein; the nucleotides encoding the C-terminal segment of YecJ downstream from the Start-Stop (orange flag and purple stop sign) were replaced with the GFP-coding sequence (placed in 0-frame relative to pTIS). The main frame stop codon is shown by the red stop sign. Right, the bar graph shows sfGFP expression efficiency as estimated by GFP/RFP/OD (%) ratio 10 h after induction of the reporter transcription. Efficiency of the reporter expression increases when the start codon of the Start-Stop site is disrupted by a mutation (construct (iTIS(-)). Mutating the stop codon of the Start-Stop site expands the length of the translated OOF internal coding sequence, which results in severe inhibition of the main frame translation. See also Figure S5 and Table S2.

Because the Start-stop sequence cannot generate any functional protein product, we surmised that the presence of the functional OOF start-stop within the main ORF may play a regulatory function, possibly affecting the efficiency of expression of the protein encoded in the main ORF. In order to test this hypothesis, we examined whether the presence of the functional start-stop affects the expression of the main ORF that hosts it. For that, we prepared a reporter construct where we fused *gfp* coding sequence to the segment of the *yecJ* ORF placing it downstream of start-stop but in frame with the *yecJ* pTIS (Figure 5D). Mutational analysis verified that expression of the YecJ-GFP fusion protein was directed by the *yecJ* pTIS (Figure 5D, wt vs. pTIS(-) bars). Notably, when the start codon of the OOF start-stop of *yecJ* was inactivated by mutation, the efficiency of expression of the YecJ-GFP reporter increased by approximately 16% (Figure 5D, wt vs. iTIS(-) bars). These results demonstrate that the presence of the active Start-Stop site within the *yecJ* gene attenuates translation of the main ORF, indicative of the possible regulatory function of the Start-Stop.

Interestingly, mutating the stop codon of the *yecJ* start-stop, that should lead to translation of a 14-codon internal ORF originating at the *yecJ* iTIS, significantly reduced the expression of the YecJ-GFP reporter (the iSTOP(-) construct in Figure 5D). This result indicates that active utilization of some of the iTISs could attenuate the expression of the main ORF whereas placing stop codon immediately after the start site of the OOF iTIS could mitigate this effect.

### Ribo-RET reveals TISs outside of the known annotated coding sequences

The ability of Ribo-RET to reveal the cryptic sites of translation initiation makes it a useful tool for identifying such unknown sites located not only within, but also outside of the annotated genes (Table S2). Thus, we have detected 6 upstream in-frame TISs (uTISs) in the *E. coli* strain BW25113 and 36 uTISs in the BL21 strain that would result in N-terminal extensions of the encoded proteins. For one gene (*potB*), we did not observe any Ribo-RET peak at the annotated pTIS (Figure S6A), suggesting that either its start site has been mis-annotated or that the annotated pTIS is activated under growth conditions different from those used in our experiments. For several other genes (e.g. *yifN*), we were able to detect Ribo-RET signals for the annotated pTIS and for the uTIS, suggesting that two protein isoforms are expressed (Figure S6B).

We also detected 41 common TISs outside of the annotated genes likely delineating the translation start sites of the unannotated short ORFs (Table S2). Some of the Ribo-RET-identified TISs define the start codons of the translated ORFs in the antisense transcripts relative to the annotated genes. Although analysis of such ORFs was beyond the scope of this study, which focused primarily on the unknown alternative internal translation start sites, the ability to detect such ORFs underscores the importance of Ribo-RET as a general tool for the genome-wide identification of translation start sites in bacteria.

## DISCUSSION

We have demonstrated the utility of the Ribo-RET approach for mapping translation initiation sites in bacterial mRNAs. Genome-wide survey of such sites in two *E. coli* strains revealed translation initiation not only at the known start codons of the annotated genes, but also at over one hundred mRNA sites nested within the currently recognized ORFs. Proteins whose synthesis is initiated as such sites may constitute a previously obscure fraction of the proteome and may play important roles in cell physiology. In addition, initiation of translation at internal codons may play a regulatory role by influencing the translation efficiency of the main protein product.

Mapping of the cryptic translation start sites was possible due to the action of RET as a highly-specific inhibitor of translation initiation, arresting ribosomes at the mRNA start codons. This finding was somewhat unexpected because binding of RET at the PTC active site would clash with the placement of formyl-methionine moiety of fMet-tRNA and by dislodging of initiator tRNA from the ribosome would prevent the start codon recognition (Gualerzi and Pon, 2015; Yan et al., 2006). However, our in vitro (toeprinting) and in vivo (Ribo-seq) data strongly argue that the RET-bound ribosome retains the P-site bound fMet-tRNA, thus allowing the antibiotic to lock the ribosome precisely at the mRNA start codons. While co-habitancy of RET and initiator fMet-tRNA in the ribosome is possible if the aminoacyl moiety of fMet-tRNA is displaced from the PTC active site, the presence of an extended nascent chain is not (Figure S1B), confining the RET inhibitory action strictly to the initiating ribosome. It is this specificity of RET action exclusively upon the initiating ribosome which makes it possible to utilize the antibiotic for confidently charting the conventional translation initiation sites at the beginning of the protein-coding sequences and also for mapping initiation-competent codons within the ORFs.

While RET readily inhibits the growth of Gram-positive bacteria it is less active against Gram-negative species, partly due to the active efflux of the drug from these cells (Jones et al., 2006). Therefore, in our experiments with Gram-negative *E. coli* we needed to use the strains that lacked the TolC component of the multi-drug transporters (Zgurskaya et al., 2011). This or similar ‘antibiotic sensitizing’ approaches could be used to apply Ribo-RET to other Gram-negative species. Even better, newer broad-spectrum, pleuromutilins, such as lefamulin, (Paukner and Riedl, 2017), could be likely used directly for mapping translation initiation sites in both Gram-positive and Gram-negative bacterial species. Other antibiotics that exclusively bind to the initiating, but not elongating ribosomes, (e.g. proline-rich antimicrobial peptides (Polikanov et al., 2018)) could also be explored for their utility in mapping translation start sites in bacteria.

Ribo-RET revealed the presence of internal start codons in over a hundred *E. coli* genes, dramatically expanding the number of putative cases of internal initiation in bacteria of which, before our work, only a few examples were known (Meydan et al., 2018). Therefore, it appears that inner-ORF initiation of translation is a much more widespread phenomenon. Previously, lactimidomycin-assisted Ribo-seq revealed initiation of translation at inner codons of many eukaryotic ORFs (Lee et al., 2012). This effect was attributed primarily to leaky scanning resulting from the poor context of the primary start site (Lee et al., 2012). Accordingly, the secondary start sites tend to cluster close to the 5’ ends of the eukaryotic ORFs. The ability of bacterial ribosomes to bind to a TIS directly, without the need to scan through the 5’ UTR, makes internal initiation within bacterial genes much more versatile because the efficiency of TIS recognition should be unaffected by its position within the gene. It is likely that at least some of these iTIS could be exploited by bacteria for expanding their proteome by generating several proteins using a single coding sequence.

Although in most cases we have little knowledge about the possible functions of the alternative polypeptides encoded in the bacterial genes, one can envision several general scenarios:

1. In-frame internal initiation generates a secondary protein whose C-terminal sequence is identical to that of the primary polypeptide. Some of the N-terminally truncated isoform can represent a protein with a specific and distinct function. One such example is the ArcB-C polypeptide expressed due to an experimentally-verified internal initiation site within the *arcB* gene.
2. If the primary protein undergoes multimerization due to interactions mediated by its C-terminal sequence, the isoform expressed from an in-frame iTIS could partake in the complex formation. Properties of such heteromers could differ from those of the homo-complexes. Several examples of heteromers formed by the protein isoforms in bacterial species have been described previously (reviewed in (Meydan et al., 2018)). Primary proteins encoded by some of the iTIS-containing *E. coli* genes (e.g. *arcB, slyB, nudF, lysU* and *wzzB*) are known to form homodimers (Maddalo et al., 2011; Moreno-Bruna et al., 2001; Onesti et al., 1995; Pena-Sandoval and Georgellis, 2010; Stenberg et al., 2005) and thus, could be candidates for the formation of heteromers with their corresponding N-terminally truncated isoforms.
3. The activity of an in-frame 5’-proximal iTIS can generate a protein similar in functions to the primary polypeptide but differing in compartmentalization. The Ribo-RET-mapped iTISs in such genes as *speA, bamA, ivy* or *yghG* are expected to direct the production of polypeptides that lack the signal sequence of the primary protein and thus may be retained in the cytoplasm. The inner-cellular functions of some of these commonly secreted proteins remain to be investigated. Similar to bacteria, some of the iTISs identified in eukaryotic mRNAs downstream from the annotated pTIS and attributed to the ‘leaky scanning’ could also alter the subcellular localization of the alternative polypeptides (Kochetov, 2008; Lee et al., 2012).
4. Because protein stability significantly depends on the nature of the N-terminal amino acid (Dougan et al., 2010), utilization of an alternative start site may generate polypeptides with varying stability that is properly tuned to the needs of the cell under specific growth conditions.
5. The utilization of the OOF iTISs generates polypeptides whose structure (and function) could be totally unrelated to those of the main protein product. Although simultaneous evolutionary optimization of two proteins encoded in overlapping but different reading frames is obviously difficult, such genes are found in many viruses (Pavesi et al., 2018). Furthermore, the available few examples of overlapping genes in bacteria (see (Meydan et al., 2018) for review), argue in favor of functionality of at least some of the products encoded in the alternative ORFs primed by the OOF iTISs.

While some of the iTISs are likely used for expanding the cellular proteome, the significance of some of the cryptic initiation sites, particularly the OOF iTISs, may lie in their regulatory role. From this standpoint, the discovery of 14 Start-Stops within the *E. coli* genes is remarkable because it is highly unlikely that the evolutionary reason for the juxtaposition of an initiator with a stop codon would be the translation of a single amino acid product. One possible function of a Start-Stop within the gene is fine-tuning the expression of the protein encoded in the host ORF. Because, in general, both initiation and termination of protein synthesis are relatively slow (Li et al., 2014; Rodnina, 2018), binding of ribosomes at the start-stop site may function as a turnstile that modulates progression of the elongating ribosomes along the ORF and, as a result, impact the protein yield. However, if such mechanism takes place indeed, it is likely subtle, because we did not observe any significant change in the ribosome density before and after the start-stop site in the Ribo-seq data collected with the untreated cells during fast-growth. Another possibility for the Start-Stop retention is that the inadvertent appearance of an OOF iTIS may lead to translation of an alternative ORF that could severely interfere with the expression of the main protein. The deleterious effect of such iTIS may be mitigated by the introduction of a stop codon immediately after the start site. In agreement with this possibility, the Start-Stop within the *yecJ* gene interferes with translation of the main ORF to a much lesser extent than translation of a more extended ORF initiated at the same iTIS (Figure 5D).

Our data indicate that Ribo-RET can be used as a reliable genome-wide tool for precise and confident mapping of the TISs in bacteria. It is conceivable, that not all of the identified iTIS directly benefit bacterial cell and a number of them could simply represent an unavoidable ‘noise’ of imprecise recognition of the start codons by the ribosome. It is also important to keep in mind that the appearance of a Ribo-RET peak shows only the *potential* of a codon to be used as a translation start site. By arresting the ribosomes at the primary start codons while allowing the elongating ribosomes to run off the mRNAs, RET treatment leads to the generation of ribosome-free mRNAs, thereby exposing putative iTISs for an easier recognition by the remaining free ribosomes. In spite of this cautionary note, several lines of evidence make us certain that many, if not all, of the Ribo-RET-identified iTISs are recognized by the ribosomes even in the untreated cells: i) for some genes with internal initiation sites (e.g. *arcB*) an increase in ribosome density downstream of the identified iTIS can be seen in the Ribo-seq data collected with the untreated cells; ii) by using reporters or directly analyzing the protein products, we have experimentally demonstrated the functionality of iTISs in several genes (e.g. *arcB, sfsA*); iii) the unique tryptic peptide corresponding to the translation product of the *gndA* gene initiated at the OOF iTIS identified by Ribo-RET has been detected by shot-gun proteomics (Yuan et al., 2018); iv) Ribo-RET readily identified iTISs in three *E. coli* genes known to contain functional internal start sites (*infB, clpB, mrcB*); v) the evolutionary conservation of the mRNA structures around some of the iTISs argue in favor of their functionality.

Several intriguing and important questions about utilization of the alternative translation start sites remained beyond the scope of the current study. While revealing the presence of alternative translation start sites in many bacterial genes, our study has not addressed the regulatory mechanisms that control the relative utilization of pTISs and iTISs. Among other factors, the interplay of the pTIS and iTIS activity could be interesting to explore. In addition, the abundance of the proteins whose translation is initiated at an iTIS is unknown. Distinguishing the secondary product initiated at an in-frame iTIS from the main protein encoded in the full ORF would be challenging for routine shot-gun proteomics, whereas identification of the proteins resulting from the activity of the OOF iTISs requires a dedicated effort, which is yet to be undertaken. More importantly, we have little knowledge of the physiological role of the alternative proteins. Even the functions of the second IF2 isoform translated from an iTIS within the *infB* gene, that has been discovered nearly half a century ago remains enigmatic (Madison et al., 2012; Miller and Wahba, 1973).

Besides iTISs, our Ribo-RET data reveal a number of the translation initiation sites outside of the annotated genes. Most of those sites delineate previously uncharacterized short genes encoded in the sense or even in the anti-sense transcripts, some of which have been previously described (Baek et al., 2017; Dornenburg et al., 2010; Thomason et al., 2015). Short proteins encoded in such ORFs may further expand the cryptic bacterial proteome (Storz et al., 2014), whereas translation of the other small ORFs could play regulatory roles (Ito and Chiba, 2013; Vázquez-Laslop and Mankin, 2014).

In conclusion, we believe that Ribo-RET presents an exceptional opportunity to map translation start sites in bacteria. Mapping the translation initiation landscape and thereby, unveiling the hidden proteome of bacteria, including the pathogenic ones, can provide us with better clues about their physiology, evolution, and virulence mechanisms.

## ACKNOWLEDGEMENTS

We thank the members of Mankin/Vazquez-Laslop laboratory for helpful discussions, G. Storz, J. Weaver (both, National Institutes of Health) and A. Buskirk (Johns Hopkins University) for suggestions about the manuscript, J.E. Barrick (University of Texas) for providing the pRXG plasmid, D. Georgellis (National Autonomous University of Mexico for advice with some experiments, Y. Polikanov and N. Aleksashin (University of Illinois at Chicago) for help with some figures. This work was supported by the grant from the National Science Foundation MCB 1615851 (to ASM and NV-L). PVB. is supported by SFI-HRB-Wellcome Trust Biomedical Research Partnership, grant no. 210692/Z/18/Z. AEF is supported by Wellcome Trust grant no. 106207.

## SUPPLEMENTARY INFORMATION

**Figure S1:**
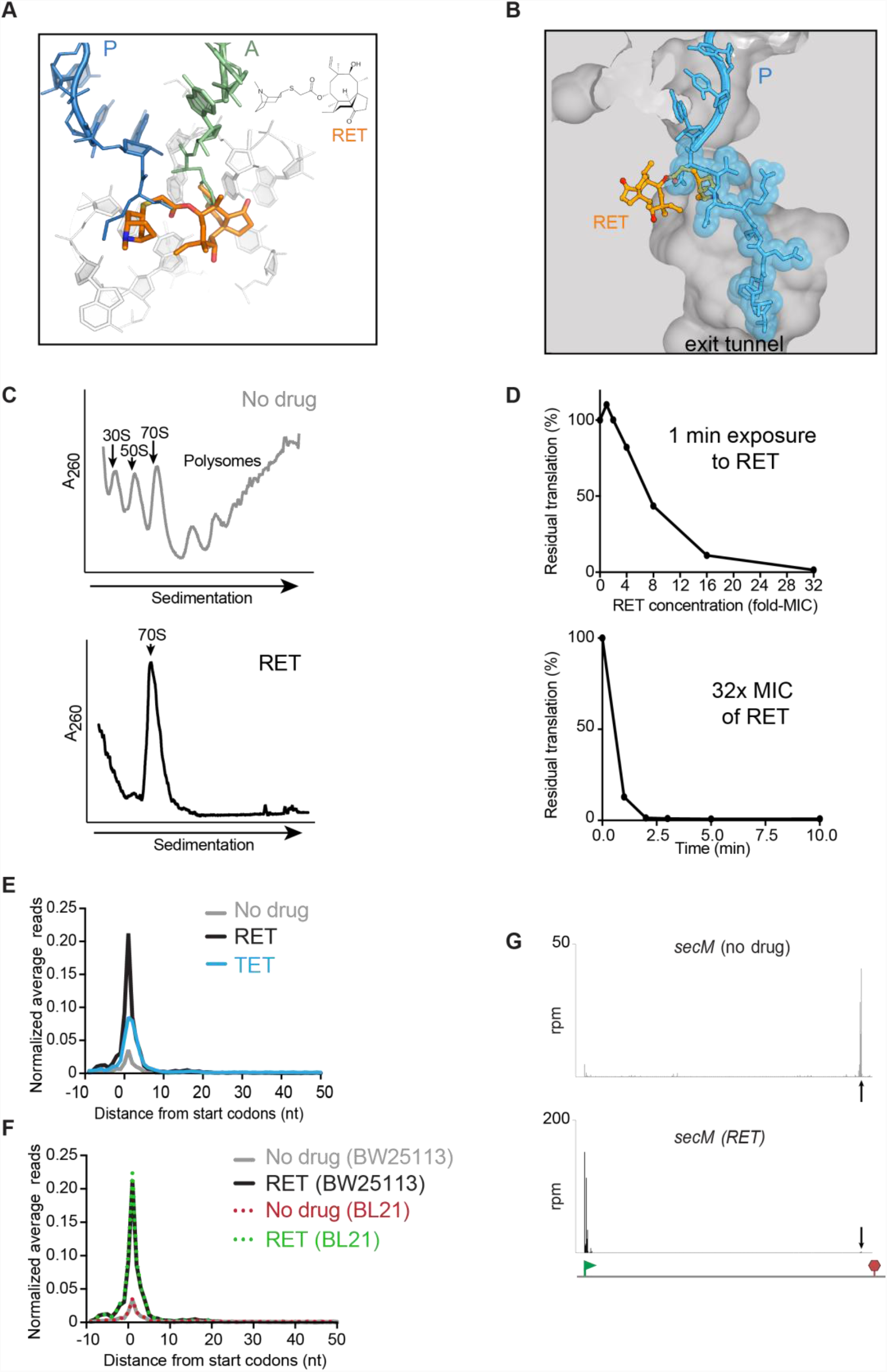
Retapamulin arrests ribosomes at initiation, Related to Figure 1. (A) The chemical structure of the pleuromutilin antibiotic retapamulin (RET) bound at the PTC active site of the bacterial ribosome. The model is based on the structural alignment of the 50S ribosomal subunit of *Deinococcus radiodurans* (*Dr*) ribosomes in complex with RET (PDB 2OGO) (Davidovich et al., 2007) and *Thermus thermophilus* 70S ribosomes with fMet-tRNA bound in the P site and Phe-tRNA in the A site (PDB 1VY4) (Polikanov et al., 2014). Note that in the 70S initiation complex, the fMet moiety of the initiator tRNA has to be displaced from the PTC active site to allow for RET binding. (B) RET cannot coexist with a nascent protein in the ribosome. Alignment of the structures of the *Dr* 50S RET complex with the *E. coli* 70S ribosome carrying ErmBL nascent peptide that esterifies P-site tRNA (PDB 5JTE) (Arenz et al., 2016). (C) Sucrose gradient analysis of polysome preparation from *E. coli* BW25113 Δ*tolC* cells untreated (top) or treated for 5 min with 12.5 µg/mL (100X MIC) RET. The shown profiles represent cryo-lyzed preparations used in Ribo-seq experiments. Qualitatively similar results have been obtained in analytical experiments with the samples prepared by freezing-thawing (see STAR Methods). (D) Residual protein synthesis in *E. coli* BL21 Δ*tolC* cells treated with RET, as estimated by incorporation of [^35^S]-methionine into the TCA-insoluble protein fraction, after 1 min exposure to increasing concentrations of RET (top) or treated with 2 µg/mL of RET (32-fold MIC) for the indicated periods of time (bottom). (E) Metagene plots comparing the normalized average relative density of ribosomal footprints in *E. coli* BW25113 Δ*tolC* cells untreated (gray trace) or treated 12.5 µg/mL (100X MIC) of RET (black trace). Blue trace represents similar analysis of the publicly-available Ribo-seq data obtained with *E. coli* BW25113 Δ*smpB* cells exposed to tetracycline (TET) [the average of two replicates of Ribo-seq experiments reported in (Nakahigashi et al., 2016)]. (F) Metagene plots comparing the normalized average relative density of ribosomal footprints in the *E. coli* strains BW25113 Δ*tolC* cells or *E. coli* BL21 Δ*tolC* untreated or treated with RET. (G) Snapshot of ribosomal footprints density in the *secM* gene of *E. coli* BW25113 ΔtolC cells untreated or treated with RET. The pTIS and stop codon of the gene are indicated by a green flag and red stop sign, respectively. The black arrow indicates the known site of translation arrest at the codon 165 of the 170-codon *secM* ORF (Nakatogawa and Ito, 2002).

**Figure S2:**
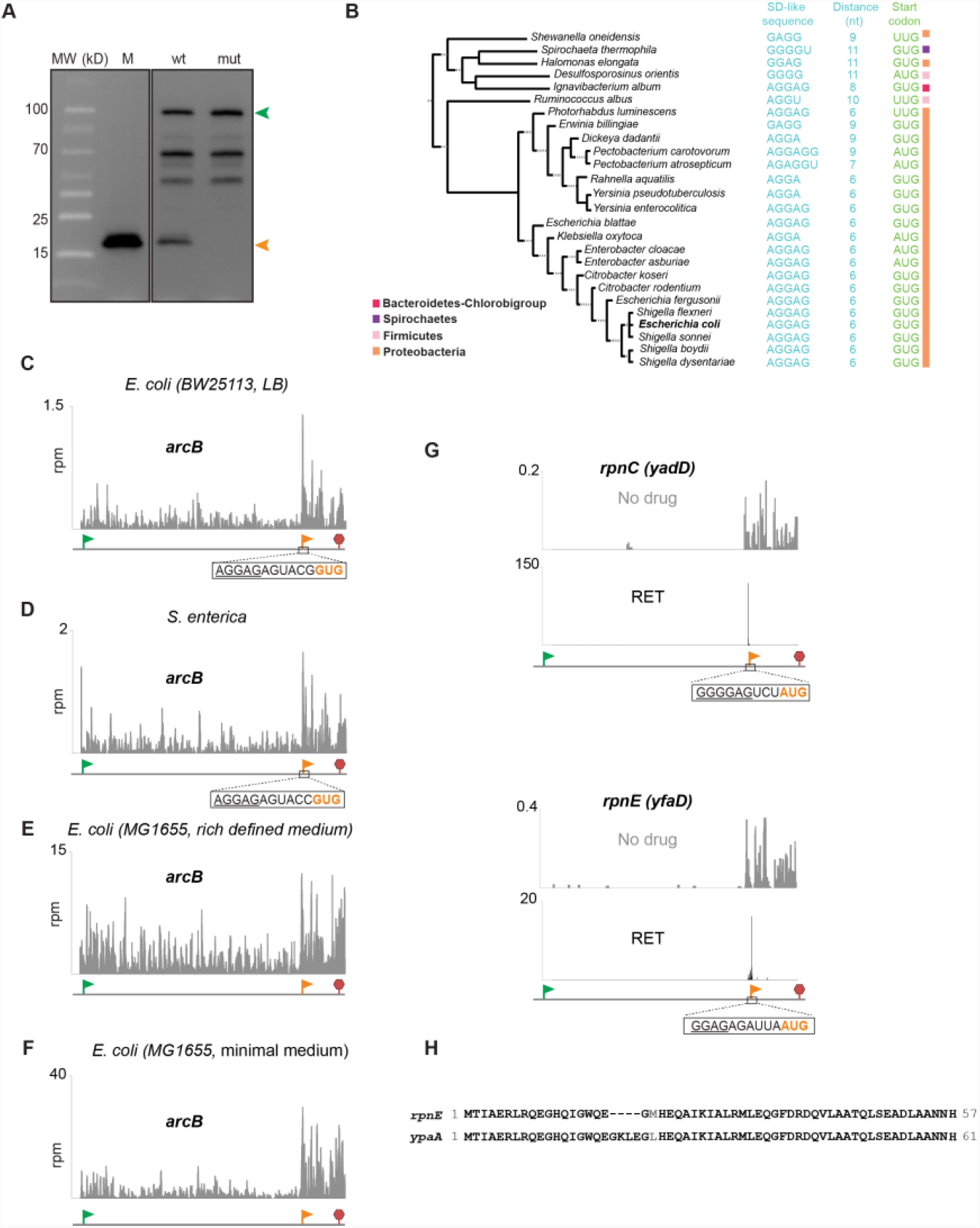
The utilization of an in-frame iTIS within the *arcB* gene leads to production of an alternative protein ArcB-C with a potential role in cell physiology, Related to Figure 3. (A) The uncropped image of the immunoblot shown in Figure 3E, representing the bands corresponding to full-length ArcB-3X FLAG and internal initiation product ArcB-C-3XFLAG (marked with arrow heads). Protein size markers are shown. The origin of the bands marked with dots is unknown. (B) The iTIS that directs translation of the ArcB-C protein is conserved in the *arcB* gene of diverse bacterial species. The putative start codons and the SD-like sequences are shown. (C-F) The upshift of ribosomal footprints in the *arcB* segment encoding ArcB-C observed in the Ribo-seq profiles of untreated *E. coli* or *Salmonella enterica* cells (Baek et al., 2017; Kannan et al., 2014; Li et al., 2014). The pTIS and iTIS of *arcB* are marked with green and orange flags, respectively, and the stop codon is indicated by a red stop sign. (G) Representative examples of Ribo-RET and Ribo-seq profiles of two out of five *coli rpn* genes. (H) Alignment of the amino acid sequence of the RpnE-C protein, translated from the iTIS within the *rpnE* gene and the protein encoded in an independent gene *ypaA*.

**Figure S3.**
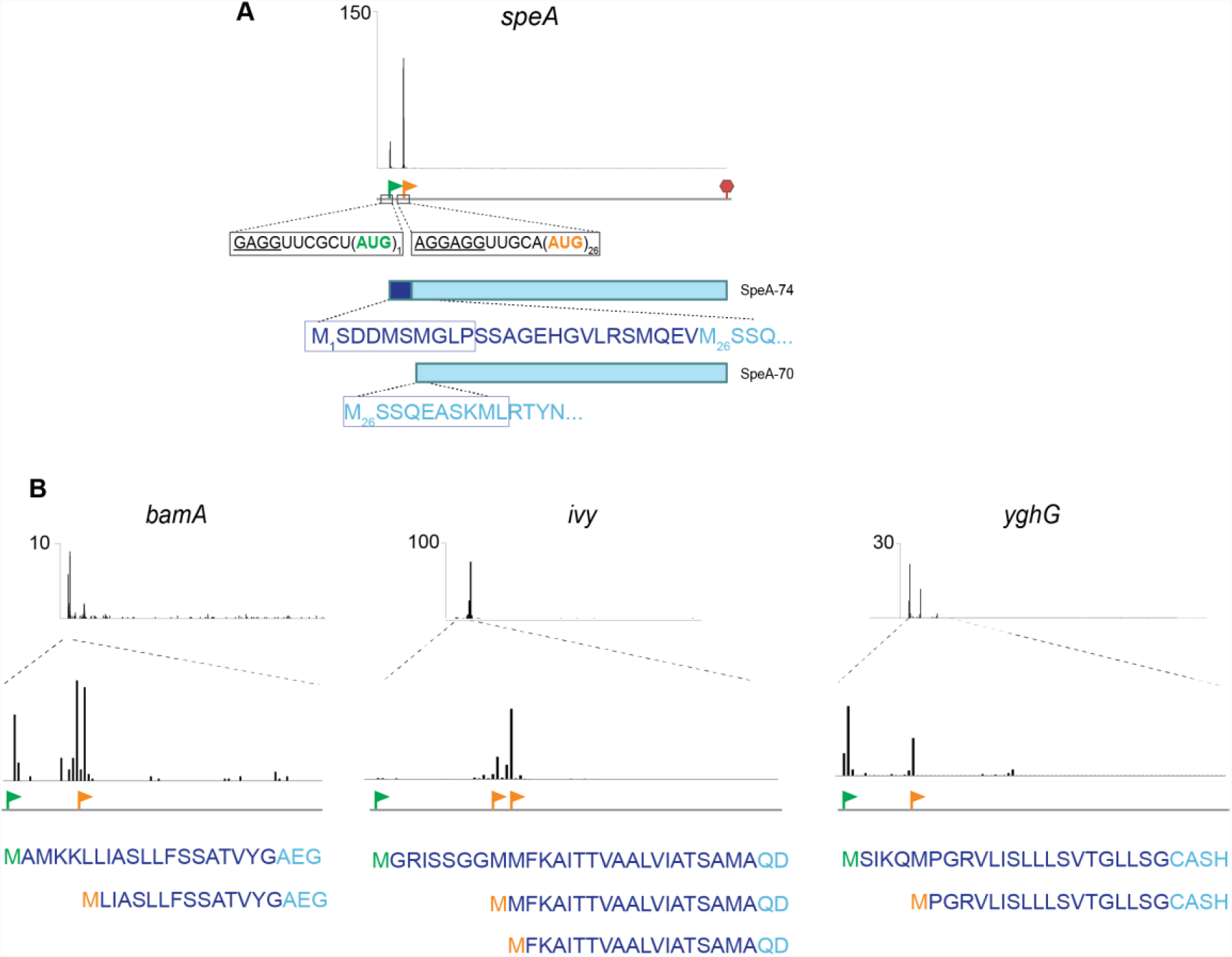
Initiation at the 5’-end proximal iTISs could produce alternative products with incomplete N-terminal signal sequences, Related to Figure 4. (A) Ribo-RET profile of the *speA* gene, showing peaks corresponding to pTIS (green flag) and iTIS (orange flag). The stop codon is indicated by a red stop sign. The putative signal sequence (indicated by dark blue letters) of SpeA-74 (Buch and Boyle, 1985) is lacking in the alternative product SpeA-70 whose translation is initiated at the iTIS. The SpeA isoforms, whose translation is initiated at the pTIS or the iTIS are expected to have different cellular localization. The peptides detected by N-terminomics are boxed (Bienvenut et al., 2015). (B) Ribo-RET profiles of *bamA, ivy* and *yghG* genes. The N-terminal amino acid sequences of the primary and predicted alternative proteins are indicated. The reported signal sequences are shown in dark blue. The pTISs of the genes are marked by green flags; iTISs are indicated with orange flags.

**Figure S4.**
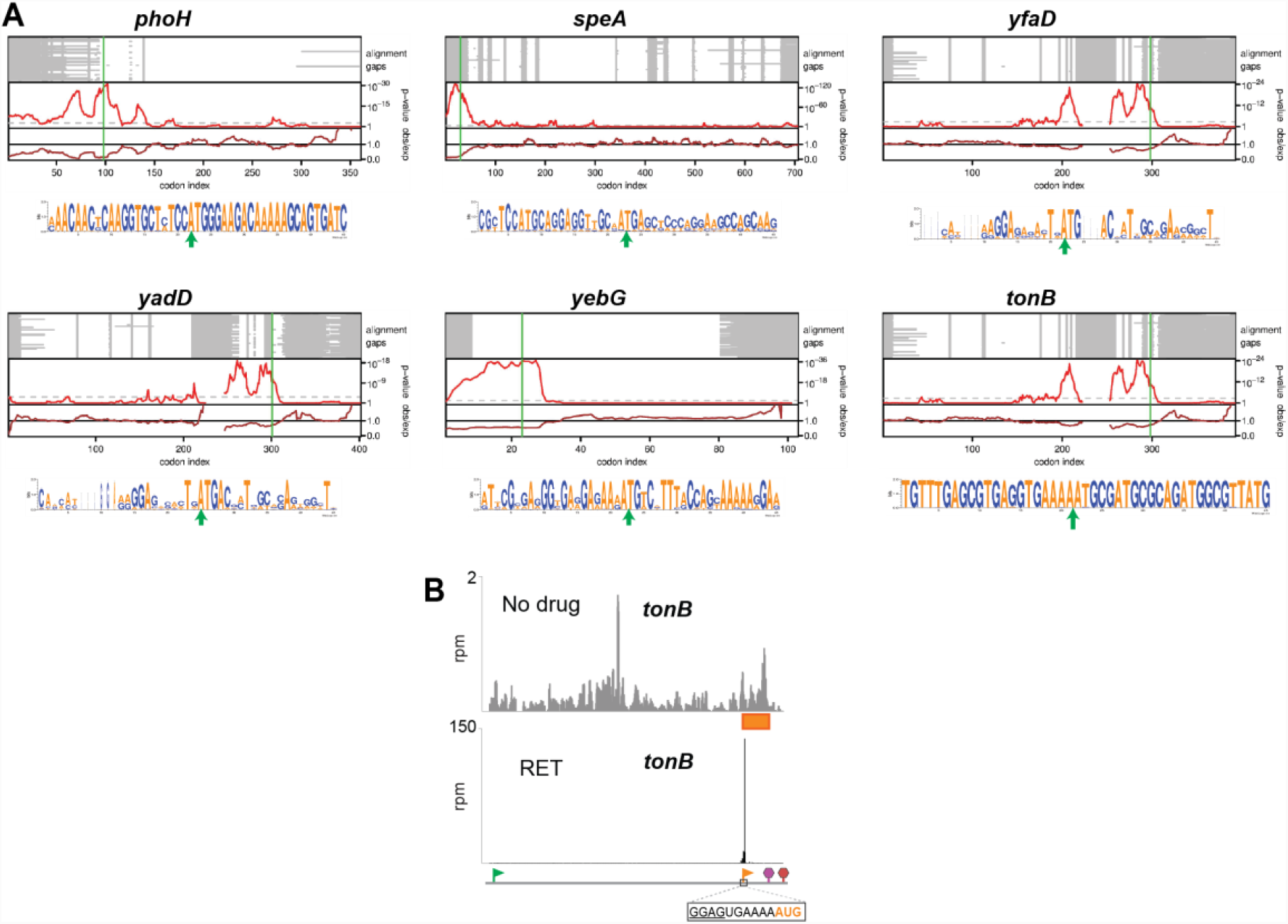
Synonymous site conservation for selected iTISs, Related to Figures 3-5. (A) Synonymous site conservation plots and weblogos for genes with in-frame iTISs (*phoH, speA, yfaD, yadD, yebG)* and for the *tonB* gene with an OOF iTIS. Alignment gaps in each sequence are indicated in grey. The two panels show the synonymous substitution rate in a 15-codon sliding window, relative to the CDS average (observed/expected; brown line) and the corresponding statistical significance (*p*-value; red line). The horizontal dashed grey line indicates a *p*-value of 0.05 / (CDS length/window size) – an approximate correction for multiple testing within a single CDS. (B) An upshift in the local density of ribosome footprints within the alternative frame defined by the *tonB* OOF iTIS (orange rectangle) in cells not exposed to antibiotic. Start codons of the pTIS and OOF iTIS are marked with green and orange flags, respectively, while the respective stop codons are indicated with red and purple stop signs. The start codon and SD-like sequence of the iTIS are shown.

**Figure S5.**
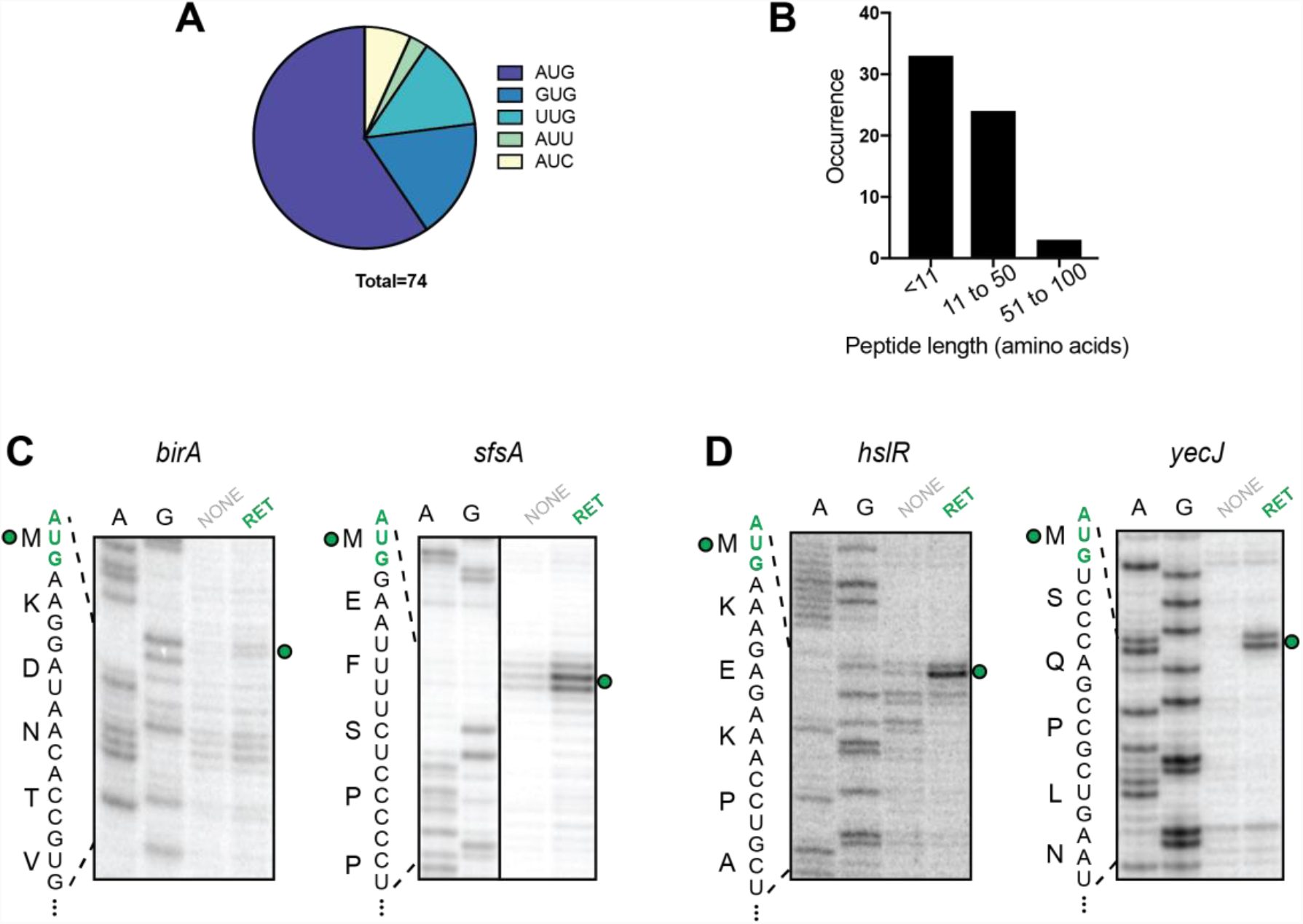
Ribo-RET reveals OOF iTISs, Related to Figures 4 and 5. (A) The distribution of start codons associated with OOF iTISs revealed by Ribo-RET. (B) The length distribution of the putative alternative proteins whose translation is initiated at OOF iTISs. (C) and (D) Toe-printing gels showing RET-induced ribosome stalling at the pTISs of *birA* and *sfsA* (shown in Figure 4) and *hslR* and *yecJ* (shown in Figure 5) genes. Samples analyzed in the lanes marked NONE contained no antibiotics. Start codons of the pTISs are indicated in green. Sequencing lanes are shown.

**Figure S6.**
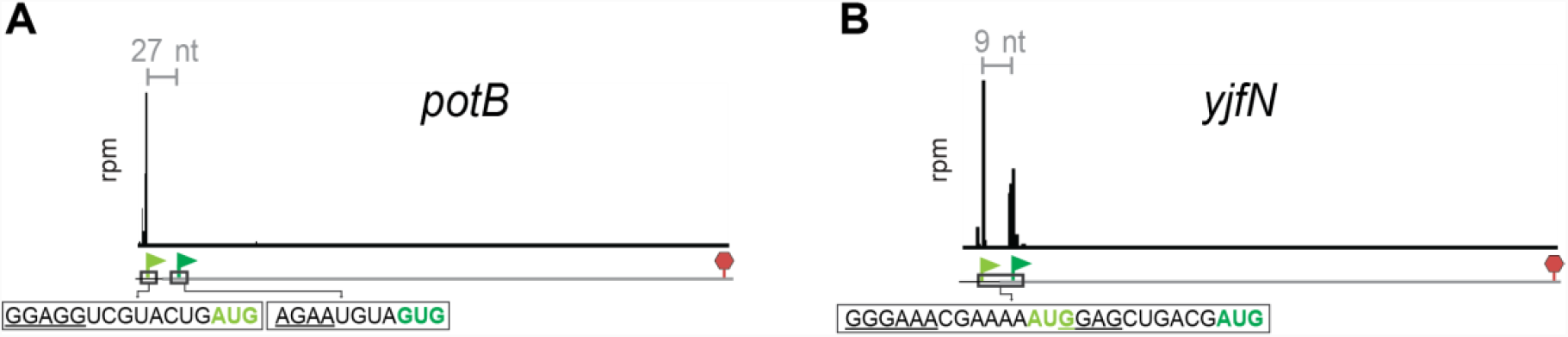
Examples of the genes with Ribo-RET identified TISs outside of the coding regions, Related to Figure 1. (A) Ribo-RET profile of the *potB* gene, shows no peak of the ribosome density at the start codon of the annotated pTIS (dark green flag), but instead reveals a strong peak at an in-frame start codon 27 nt upstream (pale green flag). (B) In the *yjfN* gene, Ribo-RET reveals peak at the annotated pTIS (dark green flag) and an additional peak 9 nts upstream from it (marked with a pale green flag). The sequences surrounding the two TISs, including the SD-like regions (underlines) are shown. See Table S2 for other cases of Ribo-RET signals outside of the coding regions.

**Scheme 1.**
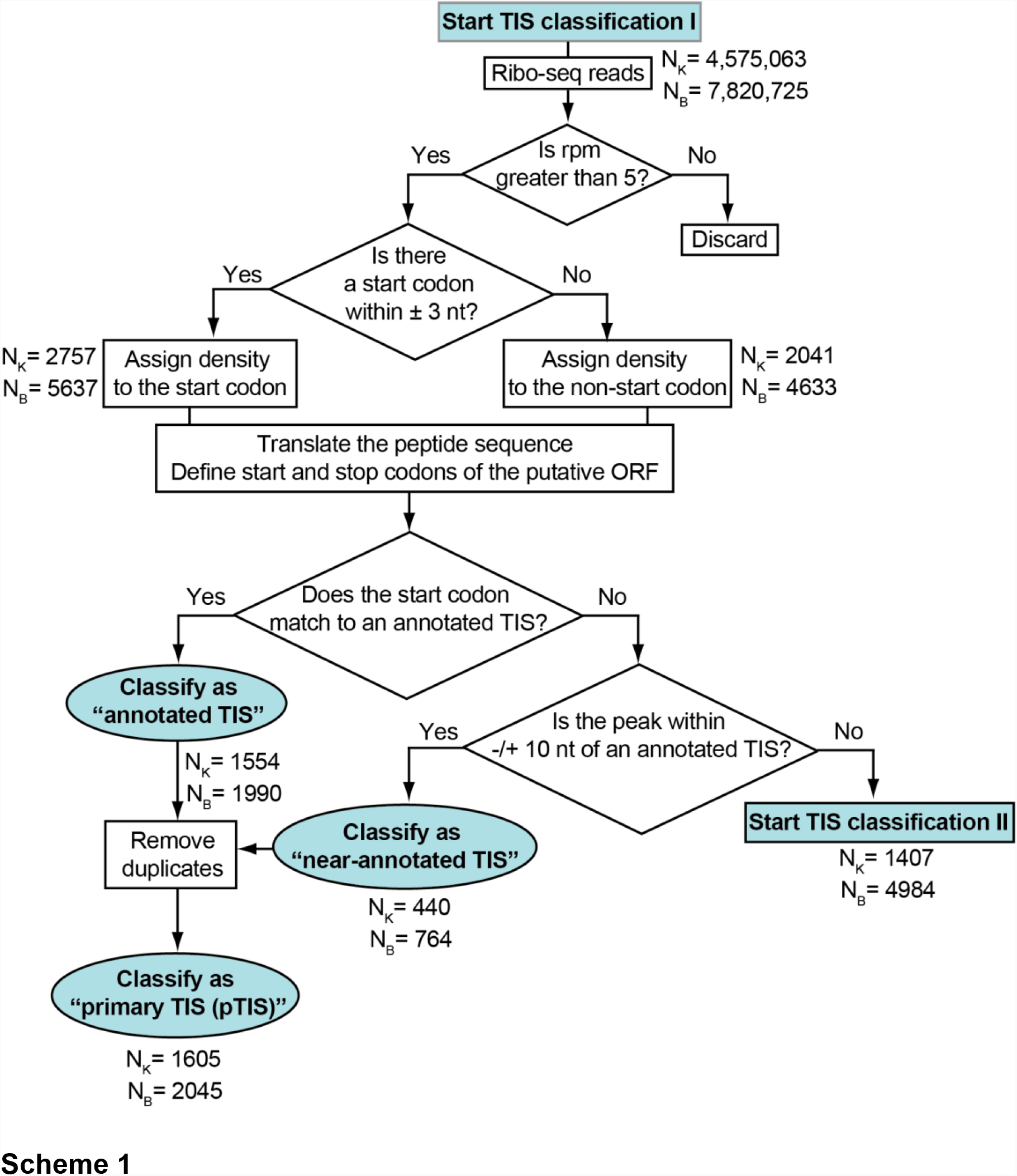

**Scheme 2.**
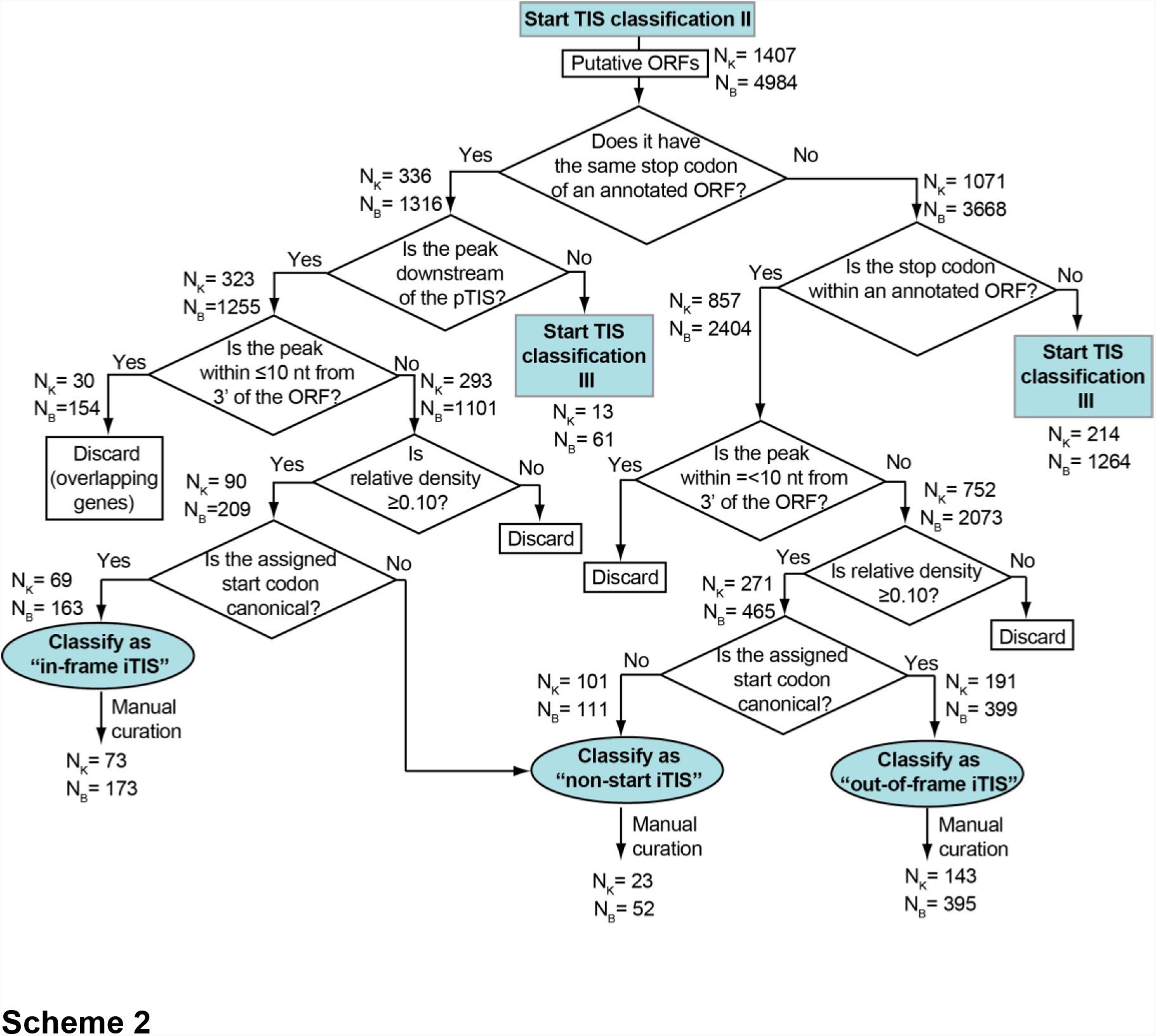

**Scheme 3.**
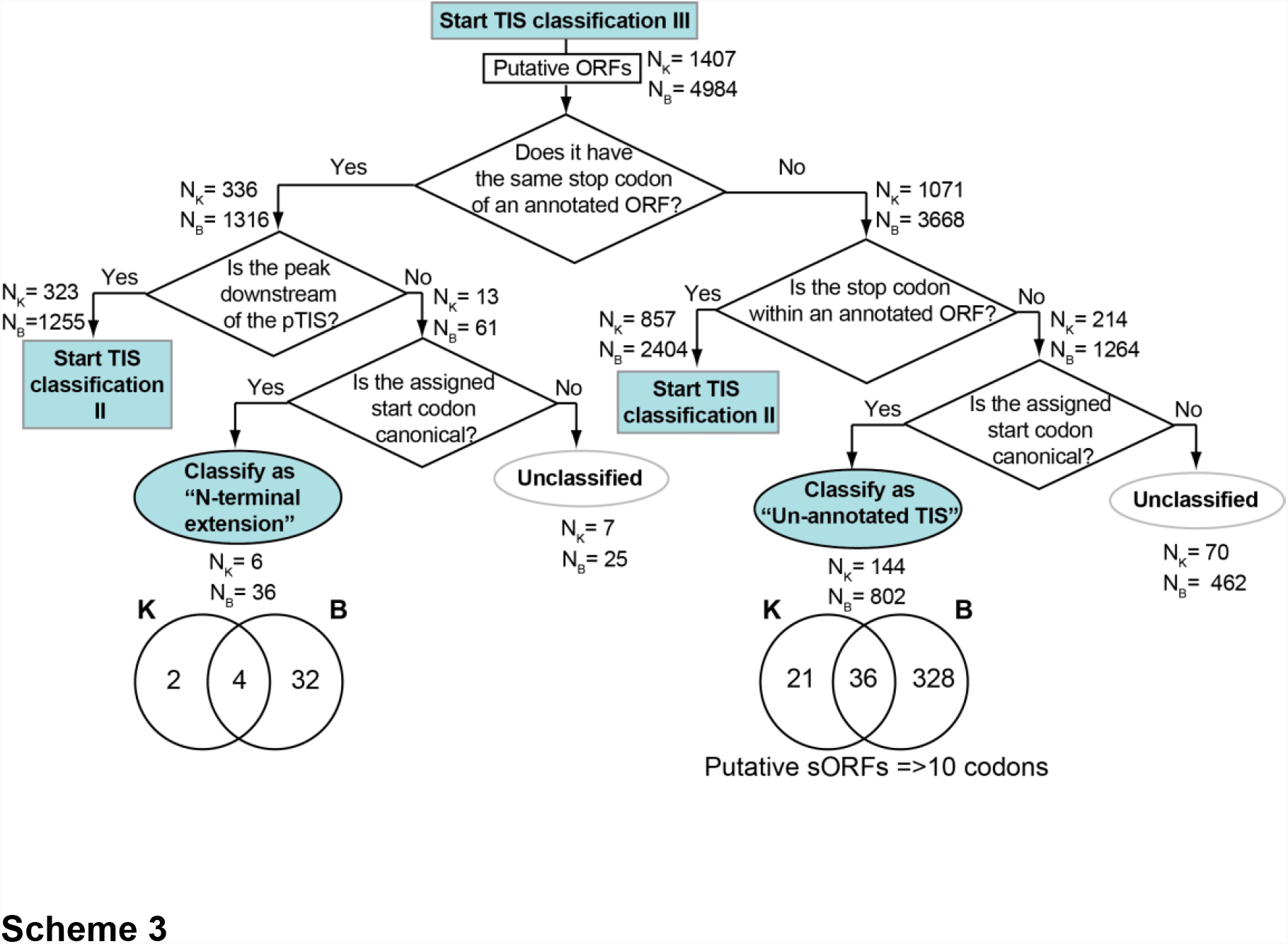

**Table S1:** List of common and strain-specific iTISs identified by Ribo-RET in the *E. coli* strains BW25113 and BL21.

**Table S2:** List of common and strain-specific TISs identified by Ribo-RET in the *E. coli* strains BW25113 and BL21 outside of the annotated genes

**Table S3:** List of primers and synthetic DNA fragments used in this study

## STAR METHODS

### Bacterial strains

Ribo-seq experiments were performed in two *E. coli* strains: the K12-type strain BW25113 (*lacI*^q^, *rrnB,* Δ*lacZ*WJ16, *hsdR514*, Δ*araBAD,* Δ*rhaBAD)* that was further rendered Δ*tolC* (called previously BWDK Kannan, 2012 #81}) and the B-type strain, BL21, (F^-^, *ompT, gal, dcm, lon, hsdS*_*B*_(r_B_^-^m_B_^-^)[*malB*^+^]_K-12_ (λ^S^) and was also rendered Δ*tolC* by recombineering (Datsenko and Wanner, 2000). For that, the kanamycin resistance cassette was PCR-amplified from BW25113 *tolC::kan* strain from the Keio collection (Baba et al., 2006) using the primers #P1 and P2 (Table S3). The PCR fragment was transformed into BL21 cells (NEB, #C2530H) carrying the Red recombinase expressing plasmid pKD46. After selection and verification of the BL21 *tolC::kan* clone, the kanamycin resistance marker was eliminated as previously described (Datsenko and Wanner, 2000). In the subsequent sections of STAR Methods we will refer to BW25113(Δ*tolC*) strain as ‘K’ strain and to BL21(Δ*tolC*) as ‘B’ strain.

Reporter plasmids were expressed in the *E. coli* strain JM109 (*end*A1, *rec*A1, *gyr*A96, *thi, hsd*R17 (r_k_^−^, m_k_^+^), *rel*A1, *sup*E44, Δ(*lac-pro*AB), [F’ *tra*D36, *pro*AB, *laq*I^q^ZΔM15]) (Promega, #P9751).

### Metabolic labeling of proteins

Inhibition of protein synthesis by RET was analyzed by metabolic labeling as described previously (Kannan et al., 2012). Specifically, the B strain cells were grown overnight at 37**°**C in M9 minimal medium supplemented with 0.003 mM thiamine and 40 μg/mL of all 19 amino acids except methionine (M9AA-Met). Cells were diluted 1:200 into fresh M9AA-Met medium and grown at 37°C until the culture density reached A_600_∼0.2. Subsequent operations were performed at 37°C. The aliquots of cell culture (28 µL) were transferred to Eppendorf tubes that contained dried-down RET (Sigma-Aldrich, #CDS023386). The final RET concentration ranged from 1x MIC to 32x MIC (0.06 μg/mL to 2 μg/mL). After incubating cells with antibiotic for 3 min, the content was transferred to another tube containing 2 µL M9AA-Met medium supplemented with 0.3 μCi of L-[^35^S]-methionine (specific activity 1,175 Ci/mmol) (MP Biomedicals). After 1 min incubation, 30 µL of 5% trichloracetic acid (TCA) was added to the cultures and this mixture was pipetted onto 35 mm 3MM paper discs (Whatman, Cat. No. 1030-025) pre-wetted with 25 µL of 5% TCA. The discs were then placed in a beaker with 500 mL 5% TCA and boiled for 5 min. TCA was discarded and this step was repeated one more time. Discs were rinsed in acetone, air-dried and placed in scintillation vials. After addition of 5 ml of scintillation cocktail (Perkin Elmer, Ultima Gold, #6013321) the amount of retained radioactivity was measured in a Scintillation Counter (Beckman, LS 6000). The data obtained from RET-treated cells were normalized to the no-drug control.

The time course of inhibition of protein synthesis by RET was monitored following essentially the same procedure except that antibiotic was added to a tube with the cells and 28 µL aliquots were withdrawn after specified time and added to tubes containing 2 µL M9AA-Met medium supplemented with 0.3 μCi of L-[^35^S]-methionine. The rest of the steps were as described above.

### Ribo-seq experiments

The Ribo-seq experiments were carried out following previously described procedures (Becker et al., 2013; Oh et al., 2011). The overnight cultures of *E. coli* grown in LB medium at 37**°**C were diluted to A_600_∼0.02 in 100 mL of fresh LB media sterilized by filtration and supplemented with 0.2% glucose. The cultures were grown at 37°C with vigorous shaking to A_600_∼0.5. RET was added to the final concentration of 100X MIC (12.5 μg/mL for the K strain or 5 μg/mL for the B strain) and incubated for 5 min (K strain) or 2 min (B strain). No antibiotic was added to the control no-drug cultures. Cells were harvested by rapid filtration, frozen in liquid nitrogen, cryo-lysed in 650 µL of buffer containing 20 mM Tris-HCl, pH 8.0, 10 mM MgCl_2_, 100 mM NH_4_Cl, 5 mM CaCl_2_, 0.4% Triton X100, 0.1% NP-40 and supplemented with 65 U RNase-free DNase I (Roche, #04716728001), 208 U SUPERase · In^Tm^ RNase inhibitor (Invitrogen, #AM2694) and GMPPNP (Sigma-Aldrich, #G0635) to the final concentration of 3 mM. After clarifying the lysate by centrifugation at 20,000 g for 10 min at 4°C samples were subjected to treatment with ∼450 U MNase (Roche, #10107921001) per 25 A_260_ of the cells for 60 min. The reactions were stopped by addition of EGTA to the final concentration of 5 mM and the monosome peak was isolated by sucrose gradient centrifugation. RNA was extracted and run on a 15% denaturing polyacrylamide gel. RNA fragments ranging in size from ∼28 to 45 nt were excised from the gel, eluted and used for library preparation as previously described (Becker et al., 2013; Oh et al., 2011). Resulting Ribo-seq data was analyzed using the GALAXY pipeline (Afgan et al., 2016; Kannan et al., 2014). The reference genome sequences U00096.3 (BW25113, ‘K’ strain) and CP001509.3 (BL21, ‘B’ strain) were used to map the Ribo-seq reads. The first position of the P-site codon was assigned by counting 15 nucleotides from the 3’ end of the Ribo-seq reads. The Ribo-seq datasets were deposited under accession number GSE1221129.

### Metagene analysis

The genes with the total rpm values ≥100 in both control and RET-treated samples were used for metagene analysis of K and B strains. The published tetracycline Ribo-seq data (Nakahigashi et al., 2016) were used to generate the corresponding metagene plot. The genes separated by less than 50 bp from the nearest neighboring gene were not included in the metagene analysis in order to avoid the ‘overlapping genes’ effects.

For every nucleotide of a gene, normalized reads were calculated by dividing reads per million (rpm) values assigned to a nucleotide by the total rpm count for the entire gene including 30 nt flanking regions. The metagene plot was generated by averaging the normalized reads for the region spanning 10 nt upstream and 50 nucleotides downstream of the first nucleotide of the start codon.

### Computational identification of translation initiation sites

The assignment of RET peaks to the start codons was performed using the algorithm provided in Supplemental Information. Specifically, we searched for a possible start codon (AUG, GUG, CUG, UUG, AUU, AUC) within 3 nucleotides upstream or downstream of the Ribo-RET peak. All other codons associated with an internal RET peak were considered as “non-start” codons (Table S1).

For assessing whether Ribo-RET peaks in K strain match the annotated start codons in the genes expressed under no-drug conditions, we calculated the percentage of genes whose rpkm values were ≥100 in the no-drug conditions and whose corresponding pTIS Ribo-RET peak values were >1 rpm. More stringent criteria were used for identification of alternative Ribo-RET peaks (rpm >5). If the Ribo-RET peak matched an annotated TIS, it was classified as pTIS (Classification I, Scheme I and Table S1). “Tailing peaks” (peaks within 10 nt downstream and upstream of the start codon) around the pTIS were considered as “near-annotated TIS” and merged with the pTISs after removing duplicates. All pTISs prior to duplicate removal are provided in Table S1 (the ‘pTISs’ tabs). The Ribo-RET peaks within coding regions were considered in Classification II and were assigned as in-frame or out-of-frame iTISs depending on the position of the likely start codon (Table S1, the ‘iTISs’ tabs). Finally, the RET peaks outside of the coding regions were considered either as N-terminal extensions or unannotated ORFs (Classification III and Table S2). The criteria for each classification are detailed in Schemes I, II and III in Supplementary Information.

### **Construction of ArcB*-*expressing plasmids**

The plasmids carrying the wt *arcB* gene or its mutant variant (G1947A, G1950A, G1959C) were generated by Gibson assembly (Gibson et al., 2009). The PCR-generated fragments covering the length of wt or mutant *arcB* genes or of the ArcB-C coding *arcB* segment were introduced into *Nco*I and *Hind*III-cut pTrc99A plasmid (Amann et al., 1988). Three PCR fragments used for the assembly of wt *arcB* plasmid were generated by using primer pairs P3/P4, P5/P6 and P7/P8 (Table S3). To construct the mutant *arcB* plasmid, the PCR fragments were generated by using primer pairs P3/P9, P7/P8 and P10/P11. The insert for the ArcB-C marker plasmid assembly was acquired as a gBlock (fragment #12 in Table S3). All the plasmids were verified by Sanger sequencing of the inserts. The plasmids were introduced in the *E. coli* BW25113 strain.

### Western blot analysis of the FLAG-tagged ArcB

The BW25113 cells carrying either wt or mutant *arcB* plasmids (or the plasmid encoding the marker ArcB-C segment of ArcB) were grown overnight at 37**°**C in LB medium supplemented with ampicillin (final concentration of 50 μg/mL). The cultures were diluted 1:100 into 5 mL LB/ampicillin medium supplemented with 0.01 mM of isopropyl-β-D-1-thiogalactopyranoside (IPTG) and grown at 37°C until culture density reached A_600_∼0.5. The cultures were harvested by centrifugation. Cells were resuspended in 300 μL of B-PER(tm) Bacterial Protein Extraction Reagent (Thermo Fisher, #78248) and centrifuged at 16,000 g for 10 min. Ten μL of the cell lysate were loaded on TGX 4-20% gradient gel (Bio-Rad, #4561096). Resolved proteins were transferred to a PVDF membrane using PVDF transfer pack (Bio-Rad, #1704156) by electroblotting (Bio-Rad Trans-Blot SD Semi-Dry Transfer Cell, 10 min at 25 V). Membrane was blocked by incubating in TBST (50 mM M Tris [pH 7.4], 150 mM NaCl, and 0.05% Tween-20) containing 5% non-fat dry milk and probed with Anti-FLAG M2-Peroxidase (Sigma-Aldrich, #A8592) and anti-GAPDH antibodies (Thermo Fisher, #MA5-15738-HRP) at 1:1000 dilution in TBST. The blot was developed using Clarity Western ECL Substrate (Bio-Rad, #170-5060) and visualized (Protein Simple, FluorChem R).

### Toeprinting assay

The DNA templates for toeprinting, were prepared by PCR amplification for the respective genes from the *E. coli* BW25113 genomic DNA. The following primer pairs were used for amplification of specific genes: *atpB*: P13/P14; *mqo*: P15/P16; *birA*: P18/P19; *hslR*: P30/P31; *yecJ*: P33/P34. Two point mutations were generated in *sfsA* in order to change the stop codon of the alternative ORF from TGA to TAG because in the PURE transcription-translation system, the termination inhibitor Api137 arrest termination at the TAG stop codon with a higher efficiency (Florin et al., 2017). This was achieved by first amplifying segments of the *sfsA* gene using pairs of primers P22/P23 and P24/P25 and then assembling the entire mutant *sfsA* sequence by mixing the PCR products together and re-amplifying using primers P26/P27. Toeprinting primer P17 was used with the *atpB* and *mqo* templates. Primers P20, P28, P32 and P36 were used for analysis of ribosome arrest at pTIS of *birA, sfsA, hslR* and *yecJ* templates, respectively. Primers P21, P29, P33 and P37 were used for the analysis of ribosome arrest at the iTISs of *birA, sfsA, hslR* and *yecJ* templates, respectively.

Transcription-translation was performed in 5 µL reactions of the PURExpress system (New England Biolabs, #E6800S) for 30 min at 37**°**C as previously described (Orelle et al., 2013). Final concentration of RET, tetracycline (Fisher scientific, #BP912-100) or Api137 (synthesized by NovoPro Biosciences, Inc.) was 50 μM. The primer extension products were resolved on 6% sequencing gels. Gels were dried, exposed overnight to phosphorimager screens and scanned on a Typhoon Trio phosphorimager (GE Healthcare).

### Polysome analysis

For the analysis of the mechanism of RET action, the overnight culture of the K strain was diluted 1:200 in 100 mL of LB medium supplemented with 0.2% glucose. The culture was grown at 37°C with vigorous shaking to A_600_ ∼0.4 at which point RET was added to the final concentration of 100X MIC (12.5 µg/mL) (control culture was left without antibiotic). After incubation for 5 min at 37°C with shaking, cultures were transferred to pre-warmed 50 mL tubes and cells were pelleted by centrifugation in a pre-warmed 37°C Beckman JA-25 rotor at 8,000 rpm for 5 min. Pellets were resuspended in 500 µL of cold lysis buffer (20mM Tris-HCl, pH7.5, 15 mM MgCl_2_), transferred to an Eppendorf tube and frozen in a dry ice/ethanol bath. Tubes were then thawed in an ice-cold water bath and 50 µL of freshly prepared lysozyme (10 mg/ml) was added. Freezing/thawing cycle was repeated two more times. Lysis was completed by addition of 15 µL of 10% sodium deoxycholate (Sigma, #D6750) and 2 µL (2U) of RQ1 RNase-free DNase (Promega, #M610A) followed by incubation on ice for 3 min. Lysates were clarified by centrifugation in a table-top centrifuge at 20,000 g for 15 min at 4°C. Three A_260_ of the lysate were loaded on 11 ml of 10%-40% sucrose gradient in buffer 20 mM Tris-HCl, pH 7.5, 10mM MgCl_2_, 100 mM NH_4_Cl_2_, 2 mM β-mercaptoethanol. Gradients were centrifuged for 2 h in a Beckman SW-41 rotor at 39,000 rpm at 4°C. Sucrose gradients were fractionated using Piston Gradient Fractionator (Biocomp).

### Construction of the reporter plasmids

The RFP/GFP plasmids were derived from the pRXG plasmid, kindly provided by Dr. Barrick (University of Texas). The vector was first reconstructed by cutting pRXG with *Eco*RI and *Sal*I and re-assembling its backbone with 2 PCR fragments amplified from pRXG using primer pairs P38/P39 (*rfp* gene) and P40/P41 (*sf-gfp* preceded by a SD sequence). The resulting plasmid (pRXGSM), had RFP ORF with downstream *Spe*I sites flanking the SD-containing *sf-*GFP ORF. To generate pRXGSM-*sfsA* plasmids, *sfsA* sequences were PCR amplified from *E. coli* BW25113 genomic DNA with primers P42/P43, for the wt gene, or P42/P44, for the mutant variant, and assembled with the *Spe*I-cut pRXGSM plasmid. To generate the pRXGSM-*yecJ* reporter plasmids, the pRXGSM was cut with *Spe*I and assembled with each of the PCR fragments generated using the following primer pairs: P45/P46 (iTIS-wt plasmid), P45/P47 (iTIS(-)); P45/P48 (pTIS-wt), P45/P50 (pTIS-iTIS(-)). pRXGSM-*yecJ*-pTIS(-) and pRXGSM-*yecJ*-pTIS-iStop(-) plasmids were generated by site directed mutagenesis of the pRXGSM-*yecJ*-pTIS-wt plasmid using primers P49 and P51, respectively.

### Fluorescence and cell density measurements

*E. coli* JM109 cells carrying the reporter plasmids were grown overnight in LB medium supplemented with 50 μg/mL kanamycin (Fisher Scientific, #BP906-5). The cultures were then diluted 1:100 into fresh LB medium supplemented with kanamycin (50 μg/mL) and grown to A_600_ ∼0.5-0.8. The cultures were diluted to the final density of A_600_ ∼0.02 in fresh LB/kanamycin (50 μg/mL) medium supplemented with 0.1 mM IPTG and 120 μL were placed in the wells of a clear flat bottom 96 well microplate (Corning, #353072). The plates were placed in a Tecan Infinite M200 PRO plate reader, where they were incubated at 37°C with orbital shaking (duration: 1000 sec; amplitude: 3 mm), and measurements of optical density (at 600 nm), ‘green fluorescence’ (excitation: 485 nm; emission: 520 nm) and ‘red fluorescence’ (excitation: 550 nm and emission: 675 nm) were acquired in real time.

### Evolutionary conservation analysis

Protein sequences of the genes of interest were extracted from Ecogene database (Zhou and Rudd, 2013). Homologs for each gene were obtained by performing a tblastn search against the nr database (Altschul et al., 1990). Briefly, the nr database was downloaded on 19/01/2018 to a local server and tblastn searches were performed for each gene (parameters -num_descriptions 1000000 - num_alignments 1000000 -evalue 0.0001). Only those tblastn hits which share a sequence identity of at least 45% with the query sequence and whose length is at least 75% of the query sequence were retained. Hits that contain in-frame stop codons were also discarded. Alignments for each gene of interest were generated by first translating the nucleotide sequences to protein sequences, aligning the protein sequences using Clustal-Omega (Sievers et al., 2011) and then back-translating the aligned protein sequence to their corresponding nucleotide sequence using T-coffee (Notredame et al., 2000). To analyze the conservation of internal start codons for each candidate gene, a 45-nucleotide region containing the internal start codon and 7 codons on either side of the start codon was extracted from each alignment. Sequence logos (Crooks et al., 2004) were built using this region of alignment and visualized to assess the conservation of the internal start codon. In order to determine if there is purifying selection at synonymous positions in the alignment, Synplot2 was used (Firth, 2014). Synplot2 was applied to each alignment (window size – 15 codons) and the resulting plots were visualized to assess the degree of synonymous site variability in the region of internal start codon.

### **Analysis of the conservation of *arcB* internal start site**

Bacterial orthologs of *arcB* were retrieved from OrtholugeDB (Whiteside et al., 2013). Coordinates of <u>histidine-containing p</u>hosphotransfer domain (HPT domain) were determined with HMMSEARCH (Wistrand and Sonnhammer, 2005) using the model 0051674 retrieved from Superfamily database version 1.74 (Wilson et al., 2009). For predicting internal initiation site, 40 nt long fragments were extracted containing 12 codons upstream of the predicted beginning of HPT domain and 1 codon downstream. Potential SD-aSD interactions were estimated by scanning the fragment with aSD 5′-ACCUCCU-3′ using program RNAhybrid (version date 3/12/2010) (Kruger and Rehmsmeier, 2006) using a ΔG threshold of -8.5. The presence of in-frame initiation codons (AUG, GUG, UUG) was checked. If initiation codon was found closer than 15 nt from SD-aSD sequence, the gene was reported to have in-frame iTIS. After removing redundancy for the strains of the same species, internal in-frame iTISs were identified for 26 bacterial species. Maximum Likelihood (ML) tree for 26 *arcB* sequences was computed using ETE command line tools by executing the command: “ete3 -w standard_fasttree -a arcB_protein_seq_in.fas -o ete_output” (Huerta-Cepas et al., 2016). Final figure (Figure S2B) was produced with function PhyloTree from ETE3 toolkit (Huerta-Cepas et al., 2016).

